# Actin dysregulation induces immune evasion via oxidative stress-activated PD-L1 in gastric cancer

**DOI:** 10.1101/2024.12.18.629227

**Authors:** Yoojeong Seo, Jinho Jang, Kyung-Pil Ko, Gengyi Zou, Yuanjian Huang, Shengzhe Zhang, Jie Zhang, Sohee Jun, Wonhong Chu, Vishwa Venkatesan, Sanjana Dhakshinamoorthy, Jae-Il Park

**Affiliations:** Department of Experimental Radiation Oncology, Division of Radiation Oncology, The University of Texas MD Anderson Cancer Center, Houston, TX 77030, USA; Graduate School of Biomedical Sciences, The University of Texas MD Anderson Cancer Center, Houston, TX 77030, USA; Program in Genetics and Epigenetics, The University of Texas MD Anderson Cancer Center, Houston, TX 77030, USA

**Author notes:** Correspondence: Jae-Il Park. These authors contributed equally (Yoojeong Seo, Jinho Jang, Kyung-Pil Ko, Gengyi Zou, Yuanjian Huang).

**Keywords:** Actin, CRACD, CRAD, KIAA1211, gastric cancer, diffuse gastric cancer, reactive oxygen species, HIF1α, PD-L1, immunotherapy

## Abstract

Diffuse gastric adenocarcinoma (DGAC) is an aggressive malignancy with limited therapeutic options, poor prognosis, and poorly understood biology. CRACD, an actin polymerization regulator, is often inactivated in gastric cancer, including DGAC. We found that genetic engineering of murine gastric organoids with *Cracd* ablation combined with *Kras* mutation and *Trp53* loss induced aberrant cell plasticity, hyperproliferation, and hypermucinosis, the features that recapitulate DGAC transcriptional signatures. Notably, CRACD inactivation remodeled the immune landscape for immune evasion through PD-L1 enrichment in tumor cells. Mechanistically, CRACD loss disrupted actin dynamics, generating reactive oxygen species that activated HIF1α, which transactivated *PD-L1*. Pharmacologic inhibition of HIF1α or PD-L1 restored immune surveillance and suppressed tumorigenesis. These findings reveal a novel role of actin homeostasis in limiting cell plasticity and immune evasion, position CRACD as a potential biomarker for stratifying patients with DGAC, and highlight HIF1α and PD-L1 as actionable therapeutic targets.

## Introduction

Gastric cancer (GC) is the fifth most common malignancy and the fourth leading cause of cancer-related mortality worldwide, accounting for over 1 million new cases and approximately 769,000 deaths in 2020^1^. Although incidence rates have declined in some regions, GC remains a critical health burden in high-risk areas such as East Asia, South America, and Eastern Europe^2^. Its aggressive nature, driven by high rates of metastasis and resistance to conventional chemotherapy, results in poor prognosis^3^. Tumor angiogenesis is crucial in GC progression and is associated with unfavorable survival outcomes^4^.

Early detection remains a significant challenge owing to the lack of symptoms in early-stage GC. Traditional biomarkers, including CEA and CA19-9, are limited in sensitivity and specificity. Recent research highlights the potential of novel biomarkers like *FGFR2*, E-cadherin, and emerging noninvasive markers, including circulating tumor cells and microRNAs, to enhance early diagnosis and treatment strategies^5–7^. GC often metastasizes to lymph nodes, peritoneum, and, in women, ovaries, and has unique genetic mutations influencing metastatic patterns. Management of advanced GC typically involves a multidisciplinary approach, including chemotherapy, targeted therapies, and immunotherapy^8–12^. Despite such advances, challenges such as therapy resistance and relapse persist^8–14^. Moreover, patient stratification for specific therapies is another hurdle to be overcome.

Hypoxia, a hallmark of the tumor microenvironment, induces significant cellular stress and promotes the production of reactive oxygen species (ROS). As tumors grow, their rapid proliferation outpaces the development of new blood vessels, leading to regions within the tumor that are poorly oxygenated. Hypoxia induces various adaptive responses in cancer cells that promote their survival, growth, and metastasis. These responses are mediated by hypoxia-inducible factors (HIFs), particularly HIF1α, which regulates gene expression involved in angiogenesis, metabolism, cell survival, and invasion. ROS are key regulators of various cellular processes, including the dynamics of the actin cytoskeleton.^15, 16^ Actin dynamics, regulated by several actin-binding proteins, control the processes of polymerization and depolymerization. This enables cancer cells to remodel their cytoskeletons in response to external signals like hypoxia and migrate through the extracellular matrix. Actin cytoskeleton remodeling is crucial for cell motility, invasion, and metastasis^17, 18^. Dysregulation of actin polymerization pathways is often observed in many types of cancer and is associated with enhanced metastatic potential and poor prognosis^19^. Further, under hypoxic conditions, disruption of the actin pathway allows cancer cells to evade immune surveillance through several mechanisms, including remodeling of the actin cytoskeleton at the immunological synapse, regulation of vasculogenic mimicry, and modulation of actin-binding proteins^20^. Emerging evidence suggests that ROS can modulate actin-binding proteins, leading to changes in their activity and affinity for actin filaments^16^. This process can disrupt the balance between actin polymerization and depolymerization, resulting in cytoskeletal reorganization that affects cell shape, motility, and invasion^21^. However, the impact of actin dysregulation on ROS and HIF pathways is unknown.

CRACD (Capping protein inhibiting Regulator of Actin Dynamics; also referred to as CRAD/KIAA1211) is a tumor suppressor that participates in maintaining epithelial cell integrity^22^. CRACD promotes actin polymerization by inhibiting capping proteins (CAPZA and CAPZB), which are negative regulators of actin polymerization^22^. This mechanism is vital for preventing uncontrolled cellular proliferation and tumorigenesis. CRACD is often inactivated in several types of cancer^23^. In colorectal cancer, the loss of CRACD function destabilizes the cadherin-catenin-actin cytoskeleton complex, thereby releasing β-catenin and hyperactivating WNT signaling. This mechanism is linked to the development of intestinal mucinous adenomas^24^. In lung adenocarcinoma, CRACD loss is associated with the emergence of neuroendocrine cell plasticity^25^. Although CRACD is also often inactivated in GC, the roles of CRACD in GC remain unexplored.

This study aimed to explore the impact of CRACD inactivation on gastric tumorigenesis. By exploiting genetically engineered organoids, transplantation, and single-cell transcriptomics, we demonstrated that CRACD functions as a gatekeeper, suppressing oxidative stress-activated immune checkpoint molecules and thereby restricting immune evasion of gastric neoplasia.

## Results

### *Cracd* KO-induced hyperplasia of gastric epithelial cells

To define the gene alteration status of CRACD in cancer, we analyzed The Cancer Genome Atlas (TCGA) datasets from cBioPortal. The *CRACD* gene is commonly mutated in GC, including mucinous stomach adenocarcinoma, signet ring cell carcinoma of the stomach, diffuse gastric adenocarcinoma, tubular adenocarcinoma, and stomach adenocarcinoma (Supplementary Fig. 1A). Immunostaining of GC tissue microarrays revealed downregulation of CRACD protein expression in GC samples but ample expression of CRACD in the gastric epithelial cells of normal stomach tissue (Supplementary Fig. 1B, C). These results indicate that CRACD is often inactivated in GC.

Having determined that CRACD is downregulated in GC (Supplementary Fig. 1), we hypothesized that CRACD loss-of-function contributes to GC tumorigenesis. To test this, we analyzed the stomach tissues of *Cracd* knockout (KO) mice (Supplementary Fig. 2A). Compared with the control (*Cracd* wild-type [WT] mouse stomach), *Cracd* KO mouse stomach displayed hyperplasia, cell hyperproliferation, increased mucin deposition, and F-actin disorganization (Supplementary Fig. 2B-F). CRACD maintains the cadherin-catenin-actin complex via F-actin polymerization as a capping protein inhibitor^26^. Here, *Cracd* KO increased β-catenin and slightly reduced E-cadherin expression in gastric glands (Supplementary Fig. 2G, H), consistent with our previous finding^26^.

We also prepared gastric organoids (GOs) by isolating epithelial cells from the stomach tissues of *Cracd* WT and KO mice and culturing those cells into GOs. Compared with WT, the *Cracd* KO GOs exhibited cell hyperproliferation (Supplementary Fig. 2I-K). Further, in a differentiation culture condition (withdrawal of Wnt3A, R-Spondin, and Noggin, key factors in maintaining the stemness of GOs), unlike WT GOs, the *Cracd* KO GOs exhibited mucin deposition in the lumen and loss of cell polarity and adhesion (Supplementary Fig. 2L-O). *Cracd* KO GOs also displayed accelerated GO growth over multiple passages (Fig. 1A-C). In addition to CRACD, receptor tyrosine kinase (RTK)-RAS and TP53 pathways are often altered in GC^27, 28^. We recently reported that RAS activation (*Kras*^G12D^) combined with TP53 inactivation (*Trp53* KO) was sufficient for the development of intestinal-type gastric tumors^29^. To define the genetic interaction of *CRACD* loss with RTK-RAS and TP53 signaling, we used genetically engineered GOs^29^. By using the Cre-loxP recombination and CRISPR-based genetic manipulation, we established GOs carrying *Kras*^G12D^ activation and *Trp53* deletion in combination with *Cracd* KO to create *<u>C</u>racd^-^*^/*–*^(C) cells; *<u>K</u>ras^G12D^ Tr<u>p</u>53^-^*^/*–*^ (KP) cells; and *<u>C</u>racd^-^*^/*–*^ *<u>K</u>ras^G12D^ Tr<u>p</u>53^-^*^/*–*^ (CKP) cells (Fig. 1D, Supplementary Fig. 2N-Q). Compared with *Trp53^-^*^/*–*^, KP and CKP GOs grew faster and expressed carcinoma embryonic antigen (CEA5), a clinical marker of gastrointestinal tract tumors (Supplementary Fig. 2R-T).

**Figure 1.**
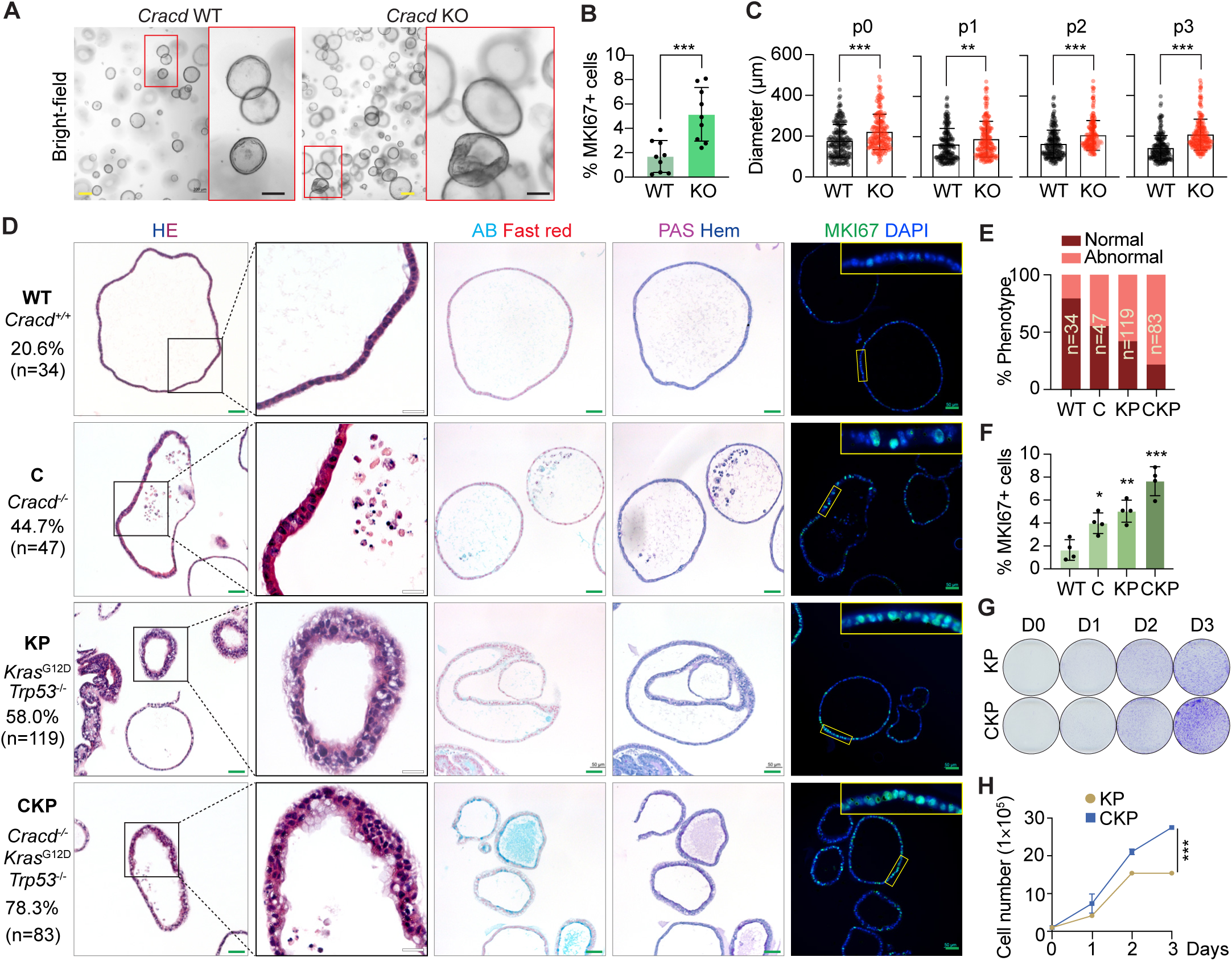
*Cracd* KO induces hyperplasia of gastric epithelial cells. **A.** Bright-field images of murine gastric organoids (GOs) (both wildtype [WT] and knockout [KO] *Cracd*); scale bars, 200 μm (yellow), 100 μm (black). **B.** Quantification of MKI67+ cells in organoids. **C.** Organoid size per number of passages. **D.** Histologic analysis of *Cracd* <u>WT</u>, *Cracd*^-/–^ (<u>C</u>), *Kras^G12D^ Trp53*^-/–^ (<u>KP</u>), and *Cracd*^-/–^ *Kras^G12D^ Trp53*^-/–^ (<u>CKP</u>) GOs by hematoxylin and eosin (HE), Alcian blue (AB), periodic acid-Schiff (PAS), and MKI6 staining. Scale bars: green, 50 μm; black, 100 μm. Percentiles indicate the ratio of abnormal morphology, such as vacuolization and cell adhesion loss, mucinous features, and cell hyperplasia. **E.** Quantification of abnormal morphology in WT, C, KP, and CKP GOs. **F.** Percentage of MKI67-positive cells in WT, C, KP, and CKP GOs. **G.** Crystal violet staining of KP and CKP cells in 2D culture (Day 0, 1, 2, and 3). **H.** Cell proliferation assay of KP and CKP cells in 2D culture. Student’s *t* test; error bars, standard deviation; n≥3. Representative images are shown. \**P*<0.05; \*\**P*<0.01; \*\*\**P*<0.001.

*Cracd* wild-type (WT) GOs develop a single layer of epithelial cells. However, C, KP, and CKP GOs displayed aberrant epithelial layer formation (Fig. 1D), with CKP GOs showing the most significant abnormal morphology, such as vacuolization and cell adhesion loss (78.3%, n=83), mucinous features, and cell hyperplasia (Fig. 1D-F). We also established GO-derived cell lines in 2D culture by using minimum growth factors^29–31^. Unlike WT and C cells, KP and CKP cells grew and maintained well at multiple passages. Consistent with the GO findings, the CKP cells showed increased colony formation and cell proliferation relative to KP cells (Fig. 1G, H). These results suggest that CRACD loss is sufficient to induce mucinous hyperplasia of the gastric epithelium.

### Cracd depletion induces cell plasticity that mirrors the transcriptional signature of gastric neoplasia

Having observed the gastric hyperplasia induced by *Cracd* KO (Fig. 1), we next investigated the cellular mechanisms provoked by *Cracd* loss. We conducted single-cell RNA sequencing (scRNA-seq) of *Cracd* WT, C, KP, and CKP GOs (Fig. 2A). Cell clusters were identified based on expression of the top 5000 genes. All datasets were integrated with the Harmony algorithm to remove the batch effect. Unsupervised cell clustering generated seven cell clusters (Fig. 2B, C). Compared with *Cracd* WT GOs (WT and KP), *Cracd* KO GOs (C [non-tumor] and CKP [tumor]) showed relatively smaller numbers of Aqp5^high^ cells (Aqp5+) and showed increased numbers of Mki67^+^ cells (proliferative) and G2/M phase cells (Fig. 2B-F, Supplementary Fig. 3A, B), implying that *Cracd* KO affects gastric epithelial cell lineages. We further used a Scissor algorithm to detect cell subpopulations linked to poor prognosis in patients with DGAC by using bulk RNA-seq datasets^32^. The proliferating cell clusters from *Cracd* KO GOs (C and CKP), which exhibited the highest similarity with bulk RNA-seq transcriptomics of DGAC (Fig. 2G), indicating that CRACD loss-of-function might be associated with DGAC tumorigenesis. Analyses of cell lineage trajectory with CytoTRACE UMAPs and pseudotime approaches showed marked alterations in cell trajectory of *Cracd* KO (WT vs. C; KP vs. CKP) (Fig. 2H, I). Whereas the Aqp5+ cell cluster served as a cellular origin in Cracd WT GOs (WT and KP), the Proliferative cell cluster (Mki67+) was the primary cellular origin of *Cracd* KO GOs (C and CKP) (Fig. 2H, I, Supplementary Fig. 3B). Likewise, consistent with the results of cell proportion analysis, the expression of *Mki67*, *Sox4*, and *Lgr5*, all key markers of cell proliferation and stem cell activity, was increased in CKP (Supplementary Fig 3A, B). In contrast, the gastric stem cell markers *Hopx*, *Lrig1*, *Prom1*, and *Aqp5* were decreased in the *Cracd* KO GOs (C and CKP) compared with WT GOs (Supplementary Fig. 3A, B). Mucinous markers, such as *Muc1*, were also increased in the *Cracd* KO organoids (C and CKP GOs) (Supplementary Fig. 3C), consistent with Alcian blue/periodic acid-Schiff staining results (Fig. 1D).

**Figure 2.**
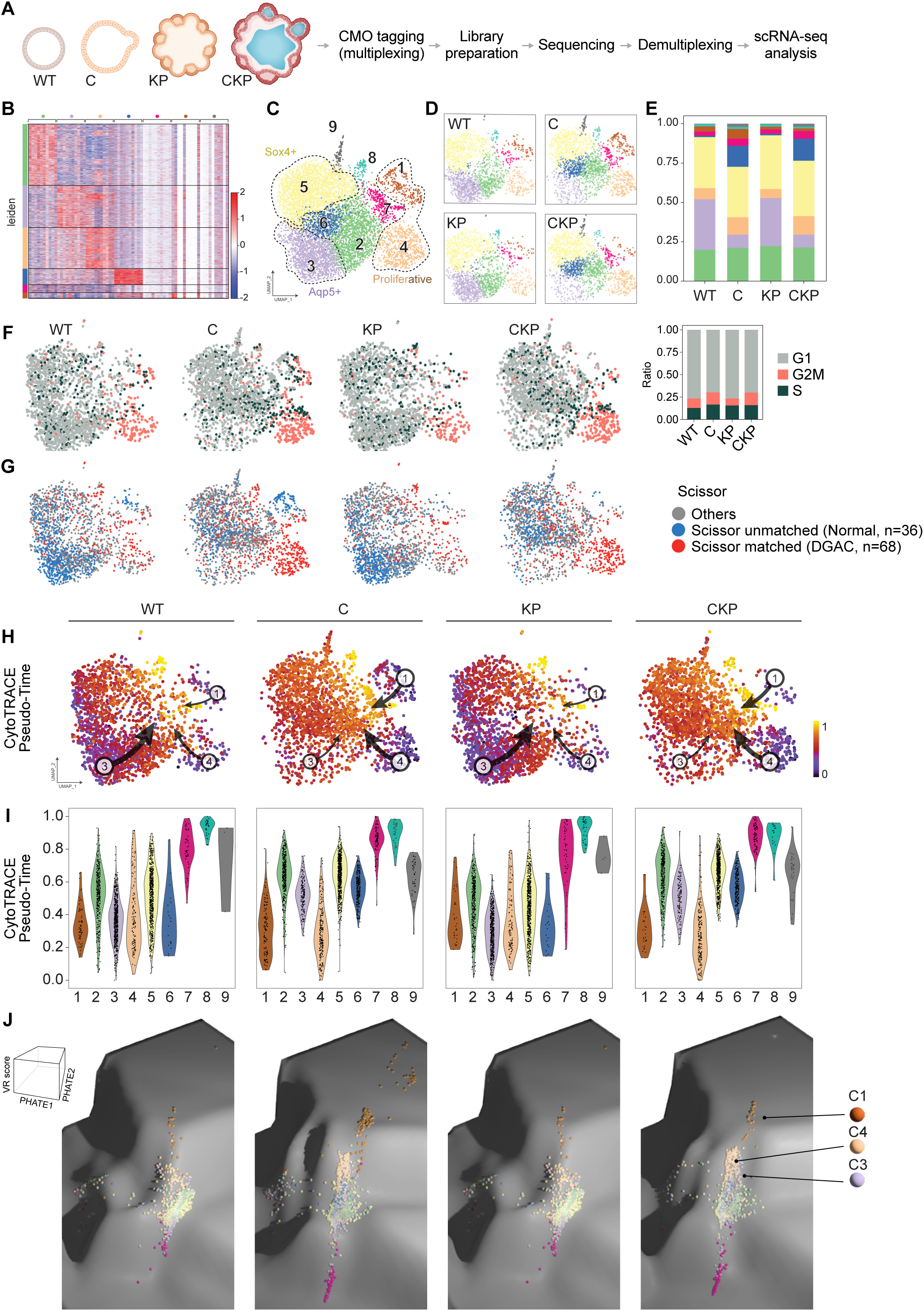
CRACD depletion induces cell plasticity of the gastric epithelia. **A.** Workflow for single-cell RNA sequencing (scRNA-seq). **B.** Heatmap displaying gene expression of markers distinguishing each cell cluster. **C.** Uniform manifold approximation and projection (UMAP) plot of cell clustering for whole cells from wildtype (WT), *Cracd* <u>WT</u>, *Cracd*^-/–^ (<u>C</u>), *Kras^G12D^ Trp53*^-/–^ (<u>KP</u>), and *Cracd*^-/–^ *Kras^G12D^ Trp53*^-/–^ (<u>CKP</u>) gastric organoids (GOs). **D.** UMAP plots showing cell clustering for WT, C, KP, and CKP GOs separately. **E.** Bar plot showing the enrichment patterns of clusters in each WT, C, KP, and CKP GO. **F.** UMAP plots depicting the cell cycle states of individual cells from WT, C, KP, and CKP GOs. **G.** UMAP plots depicting the Scissor^+^ cells from WT, C, KP, and CKP GOs. n, number of bulk RNA-seq datasets processed for Scissor analysis. **H.** CytoTRACE pseudotime inference analysis of WT, C, KP, and CKP GOs. Lower values indicate less differentiated states. **I.** Violin plots illustrating the distribution of CytoTRACE pseudotime values across each cluster for WT, C, KP, and CKP GOs. **J.** Waddington-like embedding of epithelial cells from WT, C, KP, and CKP GOs. The landscape was derived from PHATE coordinates and VR scores.

Next, we assessed the influence of CRACD depletion on cell plasticity by comparative analysis of cell plasticity potential (CPP) calculated by cell entropy, i.e., the degree of heterogeneity of gene expression and spatial information of each cell cluster visualized on PHATE maps^33^ as reconstructed by using Waddington landscape^25, 34^ (Fig. 2J). In the *Cracd* WT GOs (WT and KP), the CPP was highest in the Aqp5+ cell cluster, but in *Cracd* KO GOs (C and CKP), the CPP was higher in Proliferative cell clusters than in the Aqp5+ cell cluster (Fig. 2D, E). We also determined the pathological relevance of our models to human GC by analyzing the expression of clinical markers: *Krt7*, *Krt20*, and *Cdx2* for gastric adenocarcinoma (GAC); *Muc1*, *Muc2*, and *Muc5ac* for differentiation diagnosis; and *Sox2* and *Sox4* for undifferentiation (or stemness) markers. Dot and feature plots revealed marked increases in GAC markers in *Cracd* KO GOs (C and CKP) (Supplementary Fig. 3D, E). These results suggest that CRACD depletion induces cell plasticity that drives the transcriptional signature of gastric neoplasia.

### *Cracd* KO with *Kras*^G12D^ *Trp53* KO drives diffuse gastric tumorigenesis with immune evasion

Next, we investigated the effect of CRACD loss on gastric tumorigenesis in vivo in transplantation assays of KP and CKP cells derived from C57BL/6 mice. In immunocompromised (Nude) recipient mice subcutaneously transplanted with KP or CKP cells, the tumors derived from CKP cells were larger than the tumors from KP cells (Fig. 3A, B). We also transplanted KP or CKP cells into immunocompetent mice (C57BL/6). Unlike the KP cells in the Nude mice (100% tumor incidence), KP cells barely developed tumors in C57BL/6 mice (16.7% tumor incidence). In contrast, CKP cells generated tumors with 100% incidence in both Nude and C57BL/6 mice (Fig. 3B). These findings indicate that CKP cells can avoid immune surveillance, a feature not observed in KP cells. Moreover, unlike KP tumors, CKP tumors displayed pronounced invasiveness, penetrating the muscularis propria, increased angiogenesis and higher vascular density as well as exhibited DGAC hallmarks, including signet-ring cell carcinoma morphology (Fig. 3C-E). Histologic analysis revealed that CKP tumors were more proliferative and mucinous than KP tumors (Fig. 3F, Supplementary Fig. 6B). These findings demonstrate that CKP cells possess enhanced tumorigenic capacity, driven by increased proliferative activity and an ability to circumvent host immune defenses.

**Figure 3.**
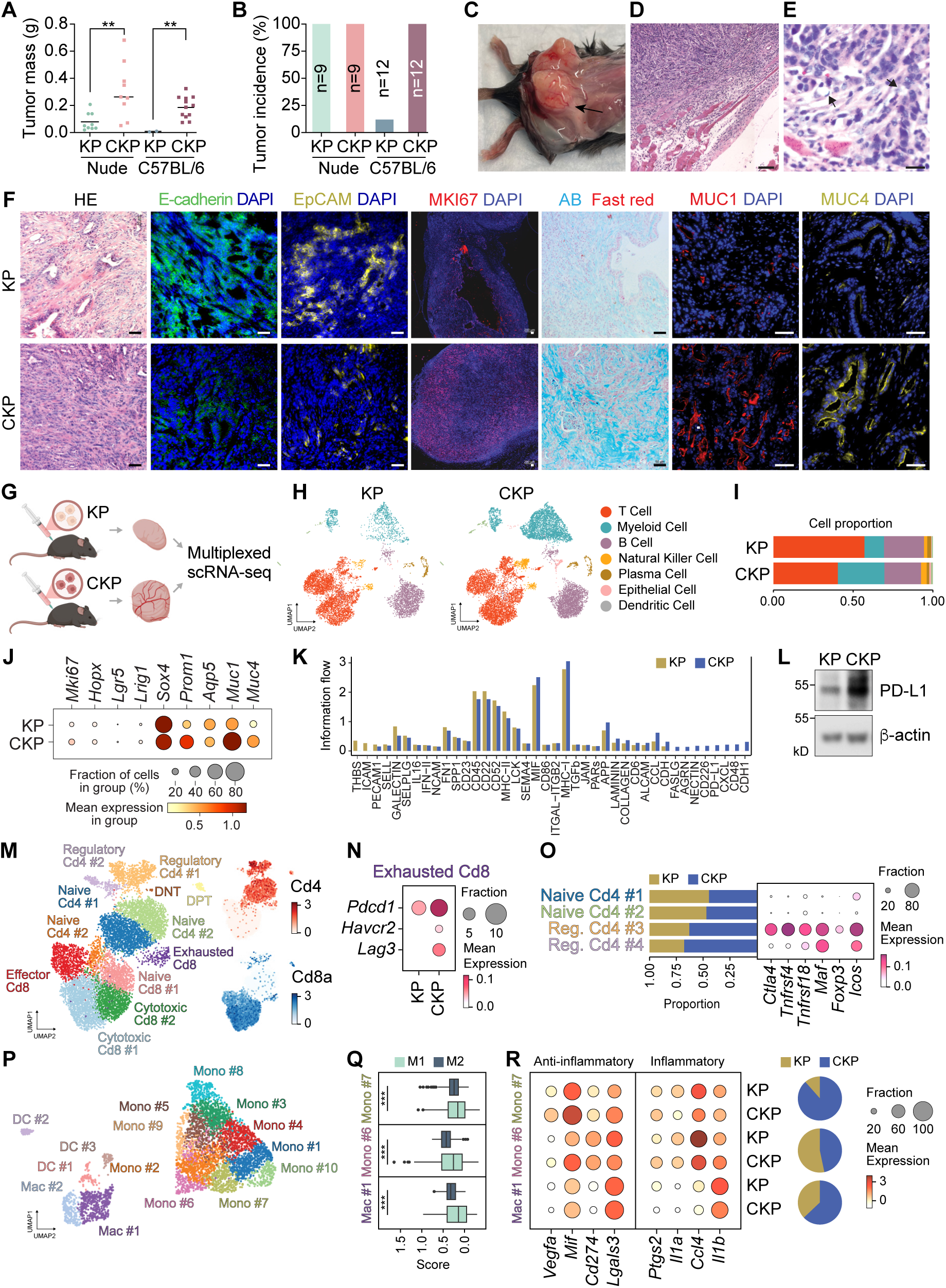
Immune evasion of *Cracd^-^*^/*–*^ *Kras*^G12D^ *Trp53^-^*^/*–*^ (CKP) tumors. **A, B.** Mass quantification (A) and tumor incidence (B) in *Kras^G12D^ Trp53*^-/–^ (KP) and CKP groups. **C.** Representative photograph of CKP allograft tumors. **D, E.** Histologic images of CKP tumors isolated from C57BL/6 mice, showing tumor invasiveness. **F.** Immunofluorescence images of KP and CKP tumors. **G.** Schematic representation of the workflow for scRNA-seq of allograft KP and CKP tumors. **H.** UMAP plots of KP and CKP tumors isolated from C57BL/6 mice. **H.** UMAP plot of cell clustering for whole cells from KP and CKP tumors. **I.** Bar plot showing the enrichment patterns of clusters in each KP and CKP tumor. **J.** Dot plot displaying the expression profiles of mucinous markers in KP and CKP tumor cells. **K.** Overall information flow of PD-L1 signaling pathway in KP and CKP tumors; CellChat analysis. **L.** Immunoblots of KP and CKP cells. Scale bars: black or white, 50 μm; yellow, 200 μm; n>3. Representative images are shown. **M.** UMAP plot illustrating the clustering of T cells from KP and CKP tumors. **N.** Dot plot displaying the expression profiles of immune checkpoint genes in the exhausted CD8^+^ T-cell subcluster (Subcluster 8) from KP and CKP tumors. **O.** Bar plot showing the enrichment patterns of T cells from KP and CKP tumors in each subcluster, accompanied by a dot plot depicting the expression profiles of Treg marker genes in CD4^+^ T-cell subclusters. **P.** UMAP plot illustrating the clustering of myeloid cells from KP and CKP tumors. **Q.** Box plots showing the M1 and M2 polarization scores in Mac #1, Mono #6, and Mono #7 subclusters, highlighting their significantly higher M2 polarization scores. **R.** Dot plot depicting the expression profiles of anti-inflammatory and inflammatory genes in Mac #1, Mono #6, and Mono #7 subcultures from KP and CKP tumors, along with their enrichment patterns across subclusters.

To determine how CRACD depletion contributes to immune evasion, we used scRNA-seq to analyze KP and CKP tumors isolated from immunocompetent mice (Fig. 3G, H). We observed a significant enrichment in myeloid cells accompanied by a depletion of T cells in CKP compared with KP, highlighting distinct differences in immune landscape between the two groups (Fig. 3H, I). In line with previous findings, CKP tumors expressed high levels of MUC1 and MUC4 compared with KP tumor cells (Fig. 3J). Next, we analyzed cell-cell interactions by using the CellChat package^32^. CKP tumors showed increased interactions mediated by PD-L1, an immune checkpoint molecule (Fig. 3K, Supplementary Fig. 4C). Because PD-L1 expression was significant in CKP tumors (Fig. 3L), PD-L1 may be a key factor driving immune evasion of CKP tumors.

To dissect the heterogeneous subpopulations of immune cells, we conducted a subclustering analysis of T cells and identified 12 distinct subclusters, each annotated based on the expression profiles of marker genes (Fig. 3M). Among these subclusters, the exhausted CD8^+^ T-cell subcluster in CKP showed marked upregulation of immune checkpoint genes, including *Pcdc1*, *Havcr2*, and *Lag3*, compared with KP (Fig. 3N). Analysis of four CD4^+^ T-cell subclusters also revealed the enrichment of two regulatory subclusters in CKP, characterized by high expression of regulatory markers such as *Ctla4*, *Tnfrsf4*, *Tnfrsf18*, *Maf*, *Foxp3*, and *Icos* (Fig. 3O).

Subclustering analysis of myeloid cells identified 15 distinct subclusters (Fig. 3P). Among those subclusters, Mac #1, Mono #6, and Mono #7 were characterized by significantly higher M2 polarization scores than M1 scores, indicating their immunosuppressive phenotypes (Fig. 3Q). Specifically, Mac #1 in CKP had decreased expression of inflammatory genes, including *Ptgs2*, *Il1a*, *Ccl4*, and *Il1b*, compared with Mac#1 in KP (Fig. 3R). In CKP, Mono #6 and Mono #7 in CKP showed upregulation of anti-inflammatory genes such as *Vegfa*, *Mif*, *Cd274*, and *Lgals3*, along with downregulation of inflammatory genes (Fig. 3R). Notably, Mono #7, which was highly enriched in CKP cells, had the most pronounced anti-inflammatory profile, implicating the immunosuppressive nature of the CKP tumor microenvironment in this phenotype (Fig. 3R). These findings suggest that CRACD loss remodels the immune landscape to allow immune evasion of tumor cells.

### CRACD depletion-elevated ROS induces HIF1α-transactivated *PD-L1*

A growing body of evidence has implicated transcriptional reprogramming in cell plasticity^35^. This led us to identify critical transcriptional modules mediating *Cracd* loss-induced cell plasticity. We identified the gene regulatory network affected by CRACD depletion by using scRNA-seq-based pySCENIC analysis^36^. The cell-regulon activity heatmap revealed distinct clusters of regulons corresponding to the specific organoid types. Clustering of regulon patterns in the UMAP space identified a dramatic shift in the gene regulatory network in *Cracd* KO cells (Fig. 4A; C [*Cracd* KO] vs. WT, CKP vs. KP). Assessment of regulon activities by Z score and regulon specificity scores both identified the activation of *Hif1a, Runx3, Sox11,* and *Sox4* regulons in *Cracd* KO (CKP) (Fig. 4B). Consistently, their expression levels (Supplementary Fig. 5A) and regulon activity levels were significantly upregulated (*P*<0.05) in *Cracd* KO compared with WT (C vs. WT, CKP vs. KP) (Fig. 4B-D, Supplementary Fig. 5A). To complement the scRNA-seq-based regulon results, we assessed the effects of *Cracd* KO on chromatin accessibility in KP and CKP cells by using an Assay for Transposase-Accessible Chromatin with sequencing (ATAC-seq) (Fig. 4E). Compared with KP control cells, CKP cells had more enriched DNA fragments binding to the transcriptional start site, increased transcription factor-binding loci, and augmented chromatin accessibility on proximal promoters (<1 kb) (Supplementary Fig. 5B, C). Specifically, the numbers of motif-predicted binding sites for Hif1a were the highest in CKP compared with KP (Fig. 4F, Supplementary Fig. 5D), indicating that *Cracd* KO cells had higher *Hif1a* activity.

**Figure 4.**
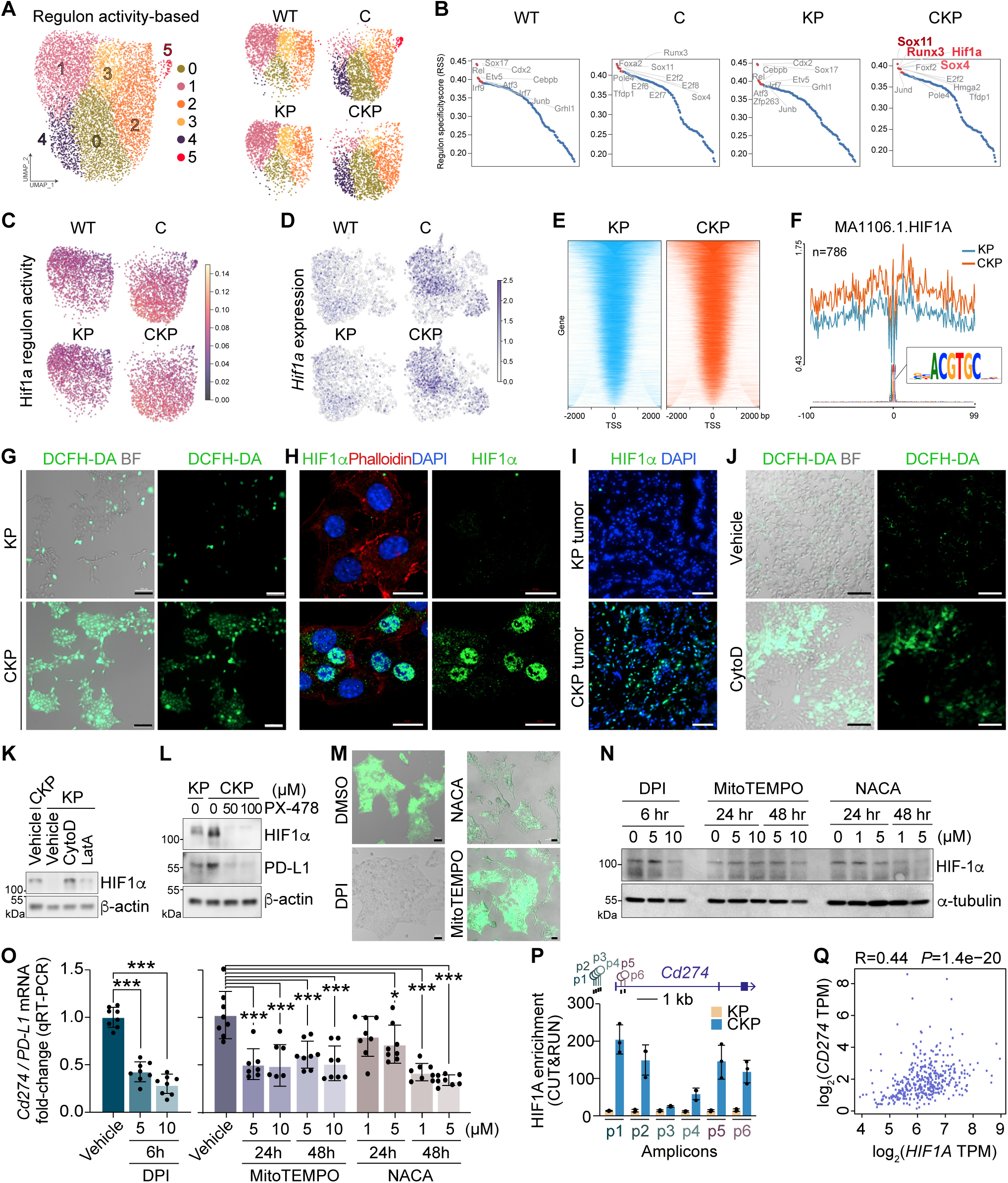
*PD-L1* transactivation by the ROS-HIF1α axis in *Cracd^-^*^/*–*^ *Kras*^G12D^ *Trp53^-^*^/*–*^ (CKP) tumor cells. **A.** Re-clustered UMAPs of organoids, annotated by unsupervised regulon patterns 0-5 (left), and separated UMAPs of organoids (right; *Cracd* wildtype [WT], *Cracd*^-/–^ [C], *Kras^G12D^ Trp53*^-/–^ [KP], and *Cracd*^-/–^ *Kras^G12D^ Trp53*^-/–^ [CKP]). **B.** Regulon specificity score plots displaying specific regulons of individual batches. **C, D.** Feature plots visualizing Hif1a’s regulon activities (C) and its mRNA expression (D). **E.** Heatmap of ATAC-seq peaks around transcription start site regions, identifying differences in gene regulation between KP and CKP. **F.** Line plot visualizing the counts of motif predicted binding sites for HIF1α (CKP vs. KP). **G.** Imaging of KP and CKP cells stained with diacetyldichlorofluorescein diacetate (DCFH-DA). Scale bars: 100 μm; BF, bright-field. **H, I.** Immunoflueorescence staining of KP and CKP cells (H) and tumors (I) for HIF1α. Scale bars: 20 μm (H) and 100 μm (I). **J.** Imaging of KP cells treated with CytoD (100 nM, 48 h) and or DCFH-DA. Scale bars: 100 μm. **K.** Immunoblot of KP cells treated with CytoD (100 nM, 48 h) or LatA (100 nM, 48 h). **L.** Immunoblot of CKP cells treated with PX-478 (72 h). **M-O.** Immunofluorescence images (M), immunoblots (N), and qRT-PCR analysis (O) of CKP cells treated with diphenyleneiodonium chloride (DPI) (5 or 10 μM, 6 h) or N-acetylcysteine (NACA) (1 or 5 μM, 48 h) or mitoTEMPO (5 or 10 μM, 48 h); scale bars, 50 μm. **P.** CUT&RUN PCR analysis of CKP cells for HIF1α. **Q.** Correlation analysis between *HIF1A* and *CD274* expression in data from The Cancer Genome Atlas (TCGA). Representative images are shown (n≥3); Student’s *t* test; error bars, standard error of the mean.

Next, we determined how CRACD loss activates the HIF1α regulon. Because HIF1α is mainly activated by ROS^37–39^, we hypothesized that actin disruption by CRACD depletion generates ROS for HIF1α activation. ROS of KP and CKP cells were visualized by 2’,7’-dichlorodihydrofluorescein diacetate (DCFH-DA) staining. Although KP cells rarely displayed ROS, CKP cells showed markedly elevated ROS (Fig. 4G). Similarly, unlike KP cells, CKP cells exhibited robust nuclear localization of HIF1α (Fig. 4H). In line with this, CKP allograft tumors displayed strong nuclear HIF1α but KP barely expressed HIF1α protein (Fig. 4I). Because CRACD loss-of-function was found to disrupt actin polymerization^23^, we tested whether actin polymerization dysfunction increases ROS. Cytochalasin D (CytoD, an actin polymerization inhibitor) was found to escalate ROS in KP cells (Fig. 4J). Similarly, CytoD and Latrunculin A (LatA, another actin polymerization inhibitor) treatment increased HIF1α protein in KP cells (Fig. 4K), suggesting that actin dysregulation generates ROS and activates HIF1α.

Several studies have shown that the HIF pathway upregulates PD-L1^40, 41^. Given the distinct expression of PD-L1 in *Cracd* KO tumors (Fig. 3K), we determined whether *Cracd* KO-upregulated PD-L1 is caused by activation of the ROS-HIF1α pathway by using inhibitors of ROS or HIF1α. Treating CKP cells with PX-478, an HIF1α inhibitor that reduces ROS, was sufficient to downregulate HIF1α and PD-L1 (Fig. 4L), indicating that the ROS-HIF1α axis mediates the upregulation of PD-L1 by loss of CRACD. To further delineate the role of ROS in HIF1α-mediated PD-L1 regulation, CKP cells were treated with ROS inhibitors, including the NADPH oxidase inhibitor diphenyleneiodonium chloride (DPI), N-acetylcysteine (NACA), and MitoTEMPO. Treatment with DPI or NACA effectively reduced ROS generation, as visualized by DCFH-DA staining (Fig. 4M). However, treatment with MitoTEMPO, a mitochondria-targeted ROS scavenger, did not significantly affect ROS or HIF1α levels (Fig. 4M, N), thereby excluding mitochondrial ROS from the CRACD loss-activated HIF1α mechanism. Intriguingly, despite the unchanged HIF1A protein levels upon MitoTEMPO treatment, we observed a reduction in *PD-L1* mRNA expression across all ROS inhibitor conditions, including MitoTEMPO (Fig. 4O). These findings suggest that although *PD-L1* transcription may depend in part on ROS-independent pathways, it remains tightly coupled to broader ROS induced by CRACD depletion (Fig. 4M-O). Next, we determined whether HIF1α directly transactivates *PD-L1* by performing Cleavage Under Targets and Release Using Nuclease (CUT&RUN) assays of KP and CKP cells. In CKP cells, HIF1α was enriched in the proximal promoters of the *Cd274/PD-L1* gene relative to the KP cells (Fig. 4P). We further examined the pathological relevance of HIF1α and PD-L1 in human GC by analyzing TCGA datasets and found that *HIF1A* mRNA expression was correlated with *Cd274/PD-L1* transcriptional expression in GC (Fig. 4Q). These findings suggest that actin dysregulation by CRACD depletion induces oxidative stress and subsequently drives the HIF1α-mediated transactivation of *PD-L1*.

### HIF1α-PD-L1 axis blockade suppresses CKP tumorigenesis

Having observed the immune evasion, along with PD-L1 enrichment, of CKP tumor cells, we evaluated the therapeutic potential of targeting the HIF1α-PD-L1 axis in CKP tumors. We first investigated the anti-tumorigenic effects of PX-478 on CKP cells. Compared with the control (vehicle), giving PX-478 to immunocompetent mice transplanted with CKP cells significantly inhibited in vivo tumorigenesis of CKP cells (Fig. 5A-C). Notably, however, PX-478 did not elicit measurable growth inhibition of KP and CKP cells in vitro (Supplementary Fig. 6A). PX-478 treatment led to a substantial increase in cleaved caspase-3 (cCas3) and a decrease in MKI67 expression (Fig. 5D-F). scRNA-seq analysis of KP and CKP tumors revealed notable reductions in T-cell populations within CKP tumors compared with KP tumors (Fig. 3H, I). Fluorescence-activated cell sorting analysis of tumor immune cells revealed marked elevations in effector T-cell populations (Granzyme B [GZMB]- or Perforin-positive) within CKP tumors after PX-478 treatment compared with control (Fig. 5G, H, Supplementary Fig. 6C-E). Given our finding that HIF1α transactivated *PD-L1* in CKP tumors (Fig. 4), we also tested whether anti-PD-L1 antibody treatment could suppress CKP tumors. Similar to PX-478, administration of anti-PD-L1 antibody markedly inhibited CKP tumorigenesis compared with IgG isotype controls (Fig. 5I, J), along with a notable increase in effector T-cell populations and tumor cell death (Fig. 5K-M), recapitulating the role of the HIF1α-PD-L1 axis in driving immune escape of CKP tumors and further indicating the tumor suppressive effect of HIF1α-PD-L1 blockade (Fig. 5N)

**Figure 5.**
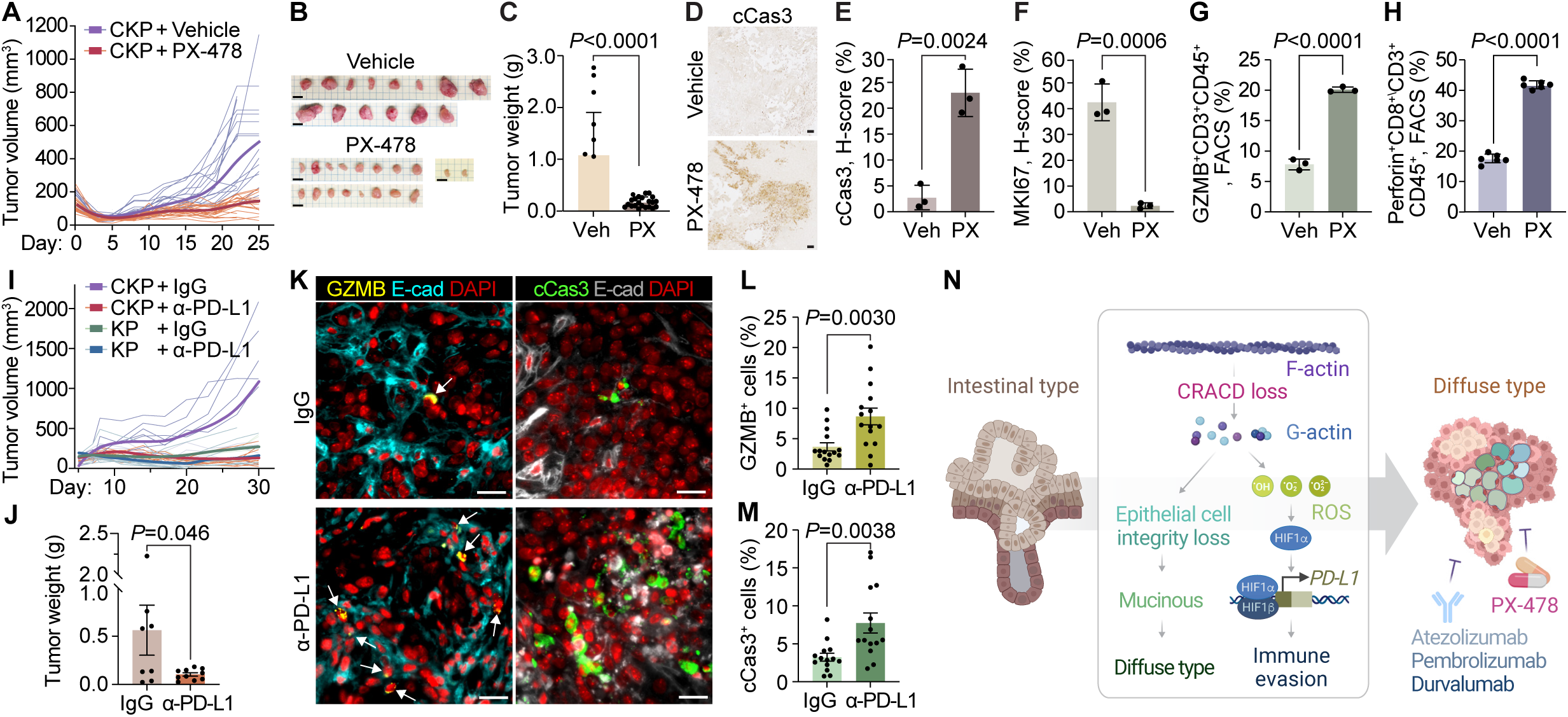
Suppression of *Cracd^-^*^/*–*^ *Kras*^G12D^ *Trp53^-^*^/*–*^ (CKP) tumorigenesis by PX-478 or anti-PD-L1 antibody. **A.** Tumor growth curves of subcutaneously transplanted CKP cells treated with vehicle (Veh) or PX-478 (PX; 40 mg/kg, by intraperitoneal injection, n=18) every other day starting on day 7-10 after transplantation. Darker lines, median values of each group. **B, C.** Images (B) and weight (C) analysis of CKP tumors harvested from vehicle and PX-478 treated groups. **D-F.** Immunohistochemical images (D) and quantification (E, F) of CKP tumors for cleaved Caspase-3 (cCas3) and MKI67. Scale bars, 100 μm. **G, H.** Florescence-activated cell sorting analysis of CKP tumors for GZMB^+^ CD3^+^ CD45^+^ (G) and Perforin^+^ CD8^+^ CD3^+^ CD45^+^ cells (H). **I, J.** Tumor growth curves (I) and weight assessment (J) of subcutaneously transplanted *Kras^G12D^ Trp53*^-/–^ (KP) and CKP cells treated with IgG or anti-PD-L1 antibody (10 mg/kg, by intraperitoneal injection). Darker lines, median values of each group. **K-M.** Immunofluorescence staining of CKP tumors for GZMB, cCas3, and E-cad (**K**) and Quantification (L, M). **N.** Illustration of gastric tumor cell plasticity and immune evasion induced by CRACD inactivation. CRACD loss of function induces cell plasticity, converting intestinal-type gastric tumors into mucinous diffuse gastric tumors. CRACD loss disrupts actin polymerization, which generates ROS. Subsequently, ROS-activated HIF1α transactivates *PD-L1* to allow immune evasion by tumor cells, which could be targeted by HIF1α inhibitors, such as PX-478 or anti-PD-L1 immune checkpoint inhibitors (atezolizumab, pembrolizumab, or duravalumab). Representative images are shown (n≥3); Student’s *t* test; error bars, SD. Panel **N** was created with BioRender.com and edited.

## Discussion

The biology underlying tumorigenesis in DGAC remains elusive. Our study revealed that CRACD is often inactivated in GC. Notably, genetic ablation of *CRACD* transformed intestinal-type GC into DGAC-like tumors characterized by hypermucinosis and immune evasion. Mechanistically, the disruption of actin polymerization from loss of CRACD led to increased ROS production, which activated the HIF1α-PD-L1 axis, driving immune evasion. Pharmacologic inhibition of HIF1α or PD-L1 restored immune surveillance and suppressed tumorigenesis, suggesting potential therapeutic strategies for CRACD-inactivated gastric tumors.

Understanding mechanisms of immune evasion is critical in cancer research, particularly as immunotherapies transform the treatment landscapes across tumor types. Tumors evolve sophisticated tactics to bypass immune surveillance, necessitating in-depth characterization of these evasion pathways to drive effective, tailored therapies. Immune evasion promotes tumor survival and underpins therapeutic resistance, underscoring the critical need to dissect its underlying biology in cancers with high immune resistance. Within this context, actin dysregulation has emerged as an area of interest. We found that actin disruption, traditionally studied for its role in cytoskeletal stability, cell migration, and epithelial integrity, affected the tumor immune landscape. We observed frequent loss-of-function in CRACD, a capping protein inhibitor that promotes actin polymerization, in GC. Unexpectedly, CRACD inactivation was found to drive immune escape through ROS activation, engaging the HIF pathway and upregulating an immune checkpoint molecule, PDL1. These findings expand our understanding of actin’s role in tumorigenesis, positioning actin dysregulation as a key factor in immune evasion with significant implications for intervention in DGAC.

Although the precise mechanisms by which actin dysregulation increases ROS remain to be elaborated, one potential pathway involves actin’s interaction with prolyl hydroxylases, enzymes central to regulating HIF1α stability. In normoxic (oxygen-rich) environments, prolyl hydroxylases hydroxylate HIF1α, marking it for degradation. However, actin may bind to and regulate prolyl hydroxylases, stabilizing HIF1α even in normoxic conditions. This mechanism would allow actin dysregulation to mimic hypoxic responses, increasing ROS and enhancing tumor-promoting signaling pathways. Further, HIF1α activation under hypoxic conditions regulates genes associated with stem-like properties and tumor progression. Therefore, actin dysregulation and HIF1α activation may synergistically promote tumor invasiveness and growth, highlighting the need to explore their interplay in DGAC.

DGAC is genetically linked to E-cadherin loss-of-function or RhoA gain-of-function^42, 43^. In this study, we considered another molecule, CRACD, loss of which was found to drive DGAC features. Interestingly, these three molecules (E-cadherin, RhoA, and CRACD) are key players in maintaining the integrity of epithelial cells by modulating cell adhesion, cell shape, cell migration, or actin cytoskeleton, implying that these mechanisms of epithelial homeostasis are pivotal in restricting aberrant cell plasticity. Moreover, additional contributors to maintaining epithelial structural resilience may be associated with DGAC tumorigenesis, thereby warranting further investigation.

In *Cracd* KO syngeneic models, we noted an increase in PD-L1-mediated immune evasion and cell proliferation. Therefore, additional mitogenic pathways activated by HIF1α are likely to be engaged in CRACD-negative GC tumorigenesis. Another intriguing observation in CKP tumors was the notable angiogenesis, represented by diffuse infiltration into the muscular layer and enriched vascular networks. This angiogenic shift likely promotes CKP tumor growth (Fig. 3C, Supplementary Fig. 6B), consistent with upregulation of vascular endothelial growth factor (*VEGF*) mRNA (Fig. 3R). Treatment with a HIF1α inhibitor effectively reduced these angiogenic and infiltrative features, suggesting that HIF1α drives the pro-angiogenic phenotype. This finding aligns with evidence that HIF1α transactivates *VEGF*, a critical driver of tumor vascularization. These data also indicate that HIF1α inhibition has therapeutic potential in CRACD-negative DGAC.

Given DGAC’s tendency toward histologic heterogeneity and poor immunotherapy response rates, our findings have significant clinical implications. The non-bulky, infiltrative nature of DGAC often complicates its early detection, and its high rates of resistance and relapse highlight the need for targeted biomarkers and treatments. CRACD could be a promising biomarker to stratify patients with DGAC, especially those with CRACD-negative GC, who may benefit from combined therapy with anti-PD-L1 antibodies and HIF1α inhibitors. At this time, cancer immunology research, despite its translational potential, is often constrained by the availability of immunocompetent preneoplastic or neoplastic model systems. To address this limitation, we established murine cell lines and GOs that recapitulate human DGAC. In our mouse allograft model, *Cracd* KO cell lines developed distinct DGAC traits, including diffuse, infiltrative growth beneath the skin and larger tumor volumes compared with KP tumors (Fig. 3). Single-cell transcriptomics also showed the similarity of transcriptional signatures between CKP organoids and DGAC tumor cells (Fig. 2G). These models offer a robust platform for investigating immune interactions and testing therapeutic strategies in DGAC.

DGAC is highly metastatic relative to intestinal-type GC^44^. To elucidate CRACD’s contribution to the metastatic potential of DGAC, further studies should examine the effects of CRACD loss on metastasis and invasion-related pathways. Additional research could clarify whether CRACD inactivation directly enhances DGAC metastasis, providing insights into mechanisms of metastasis and supporting CRACD as a potential target to limit or prevent cancer spread. Nonetheless, given our study’s reliance on murine models, translating these findings to human pathology remains challenging. Larger scRNA-seq datasets could facilitate the stratification of CRACD-positive vs. CRACD-negative GC, enhancing the relevance of our results.

GC remains a formidable malignancy owing to limited therapeutic options and poor outcomes with current immunotherapies. Our findings highlight the potential of biomarker-guided therapies for patients with CRACD-negative disease or those with DGAC. HIF1α inhibitors and anti-PD-L1 immune checkpoint blockade could provide promising therapeutic options for such patients. Furthermore, our study has established a preclinical model that effectively mimics human DGAC, offering valuable tools for drug screening and further research into GC immunology.

## Author contributions

Y.S.: Methodology, investigation, analysis, data curation, writing (original draft, review, and editing), visualization; J.J.: Methodology, investigation, software, analysis, data curation, writing (original draft, review, and editing), visualization; K.-P.K.: Methodology, investigation, software, analysis, data curation, writing (original draft), visualization; G.Z.: Methodology, investigation, analysis, data curation, writing (original draft), visualization; Y.H.: Investigation, software, analysis, writing (original draft); S. Z.: Investigation, analysis; J.Z.: Investigation; S.J.: Investigation; W.C.: Investigation; V.V.: writing (original draft); S.D.: writing (original draft); J.-I.P.: Conceptualization, methodology, analysis, writing (original draft, review, and editing), visualization, supervision, project administration, funding acquisition

## Acknowledgments

We thank Joel M. Sederstrom (Baylor College of Medicine) for technical assistance and Christine F. Wogan (Division of Radiation Oncology, MD Anderson) for manuscript editing. This work was supported by the Cancer Prevention and Research Institute of Texas (CPRIT) (RP200315 to J.-I.P.) and the National Cancer Institute (K99 CA286761 to K.-P.K., R01 CA193297, R03 CA256207, R03 CA278967, R01 CA278971, and R01 CA279867 to J.-I.P.). The core facilities at MD Anderson (DNA sequencing and Genetically Engineered Mouse Facility) were supported by the National Cancer Institute Cancer Center Support Grant (P30 CA016672). The core facilities at Baylor College of Medicine (Cytometry and Cell Sorting Core and Single Cell Genomics Core) were supported by CPRIT (RP180672 and RP200504) and the National Institutes of Health (CA125123, RR024574, S10OD023469, S10OD025240, and P30EY002520).

---

**Supplementary Figure 1.**
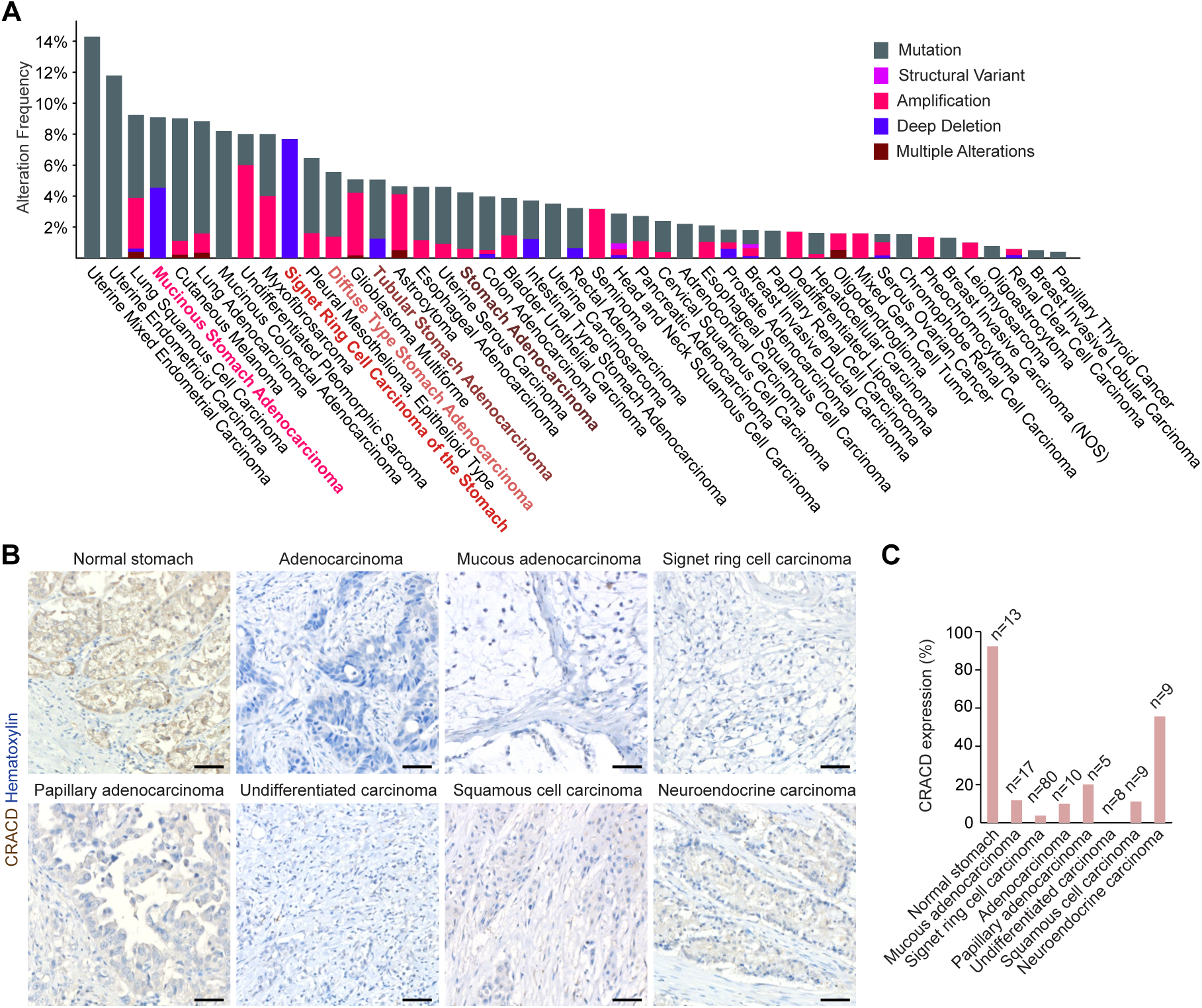

**Supplementary Figure 2.**
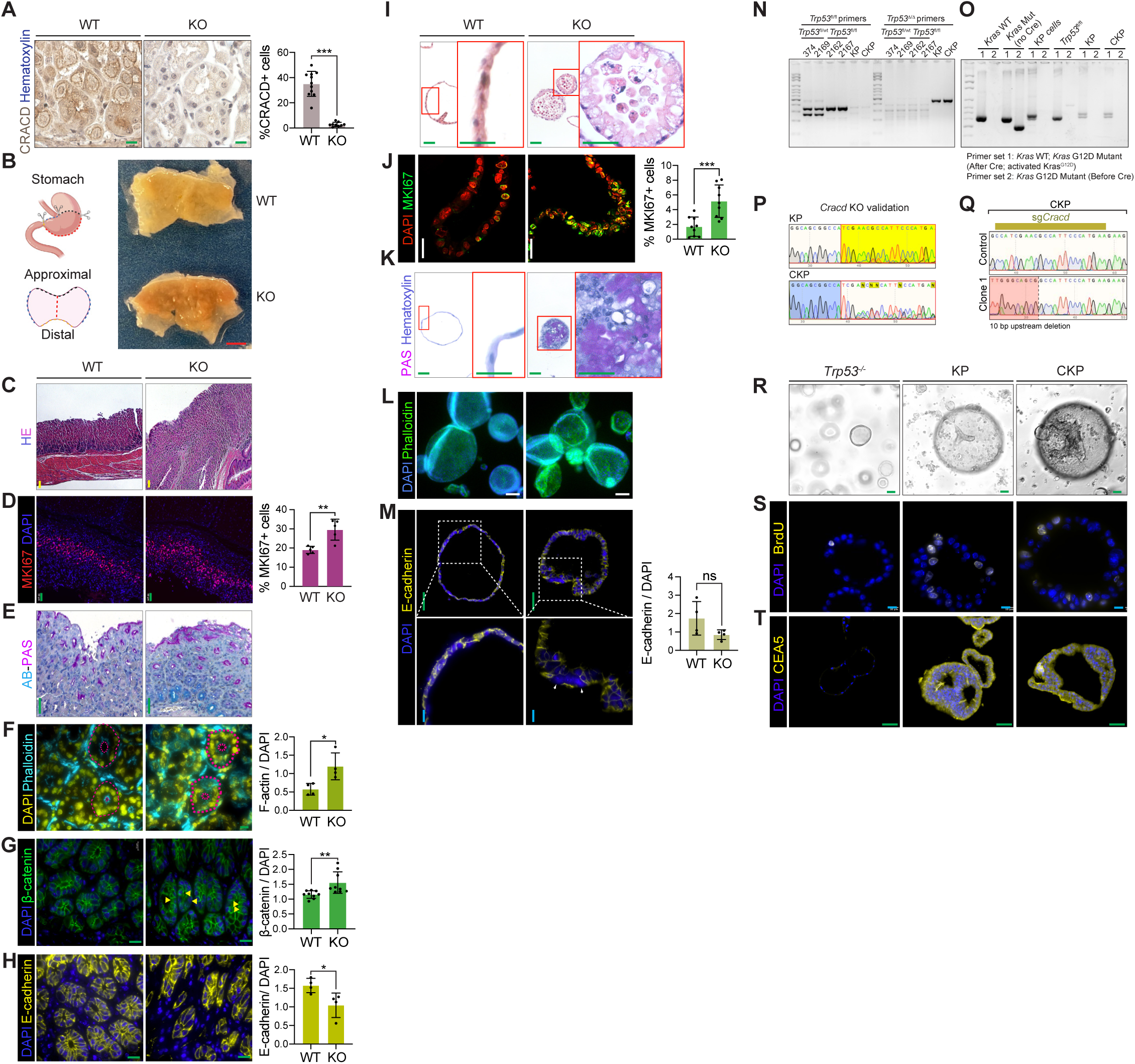

**Supplementary Figure 3.**
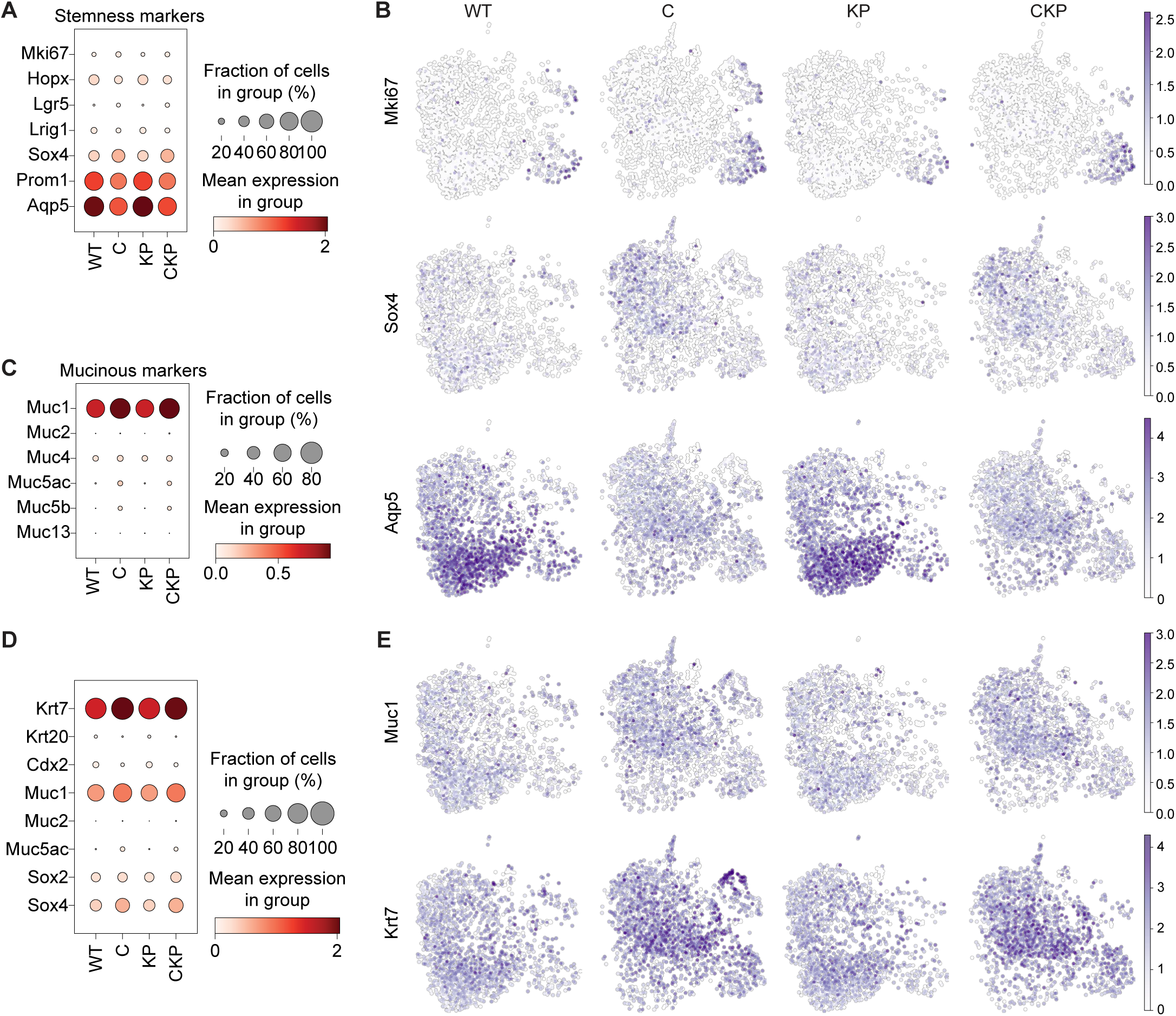

**Supplementary Figure 4.**
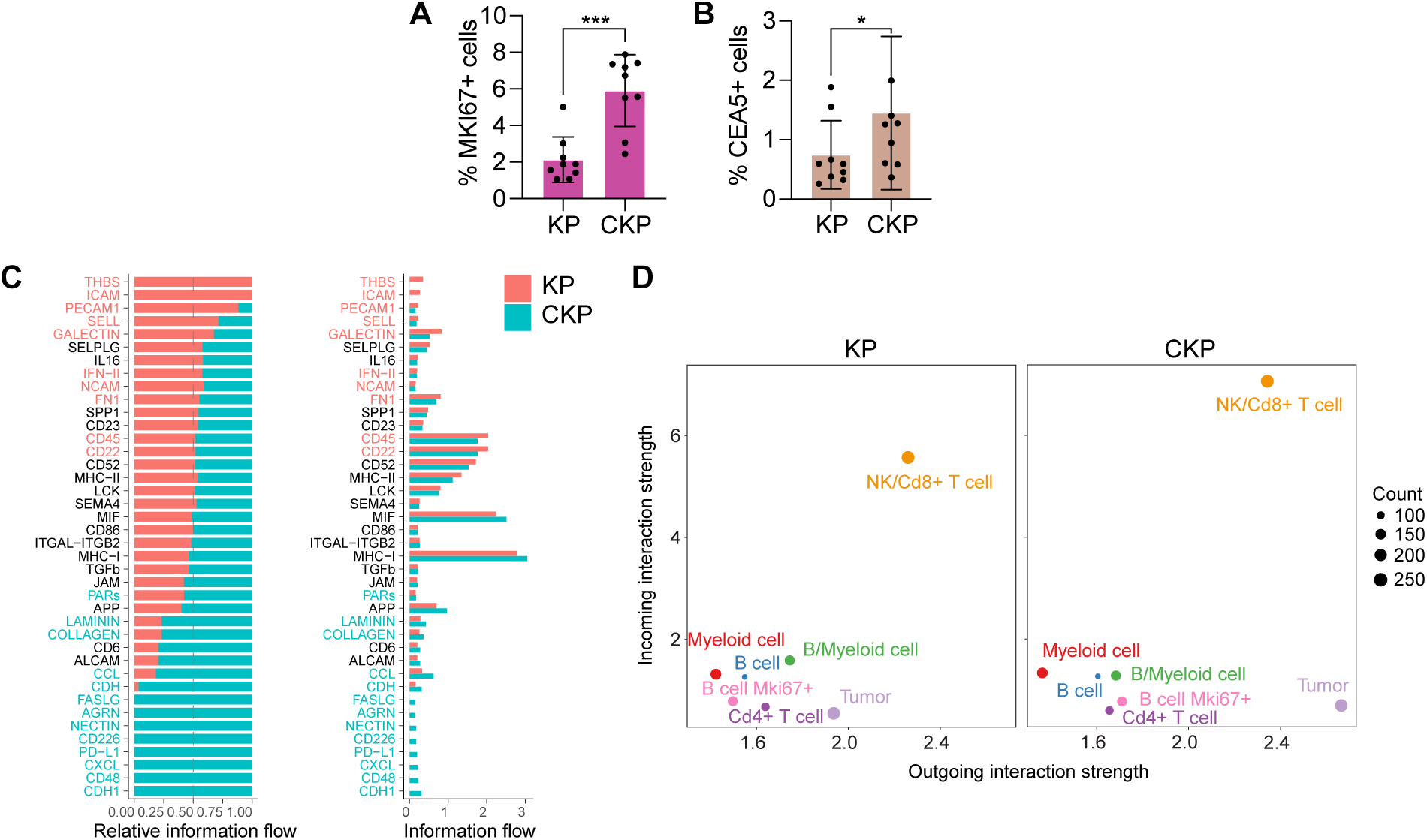

**Supplementary Figure 5.**
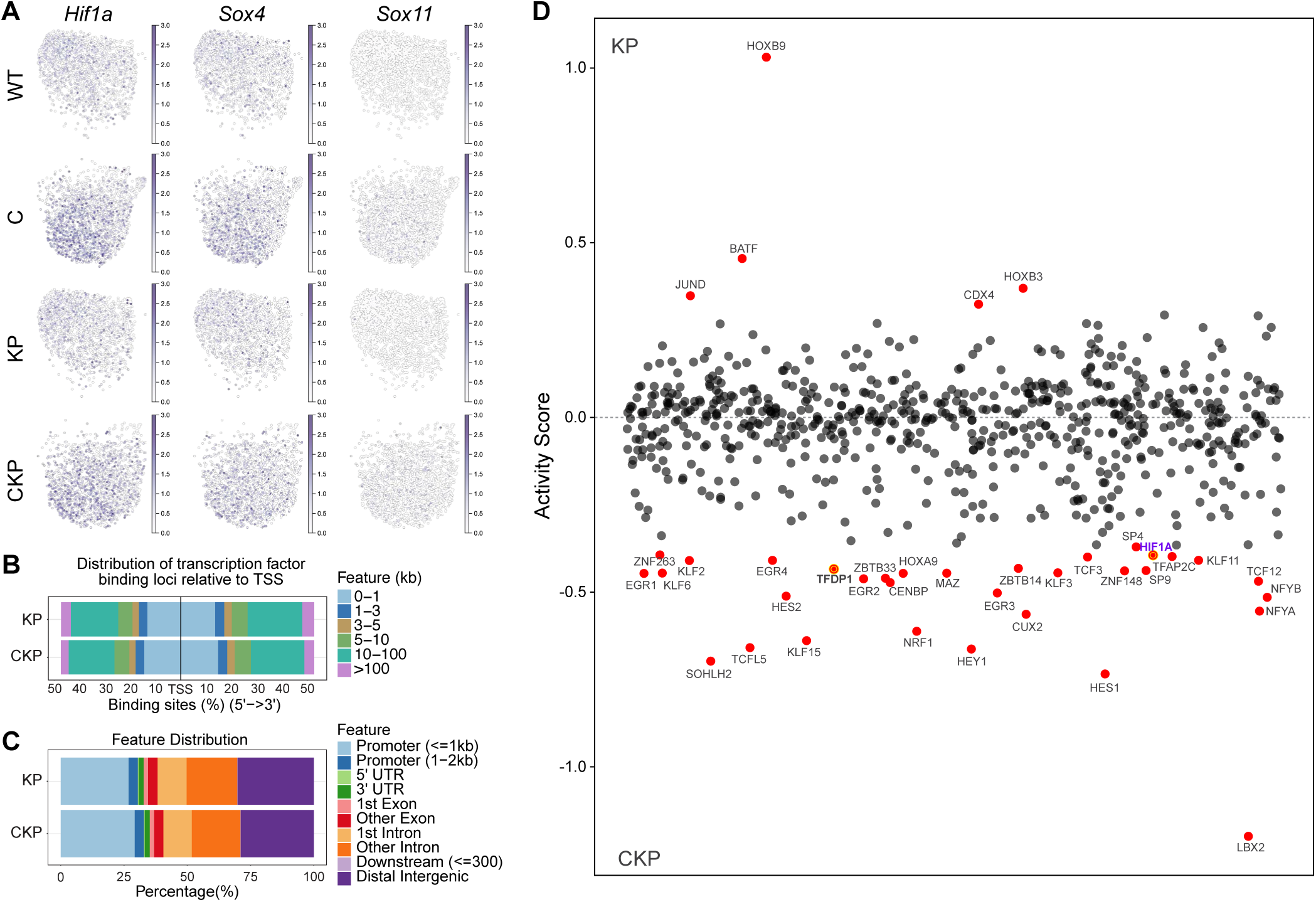

**Supplementary Figure 6.**
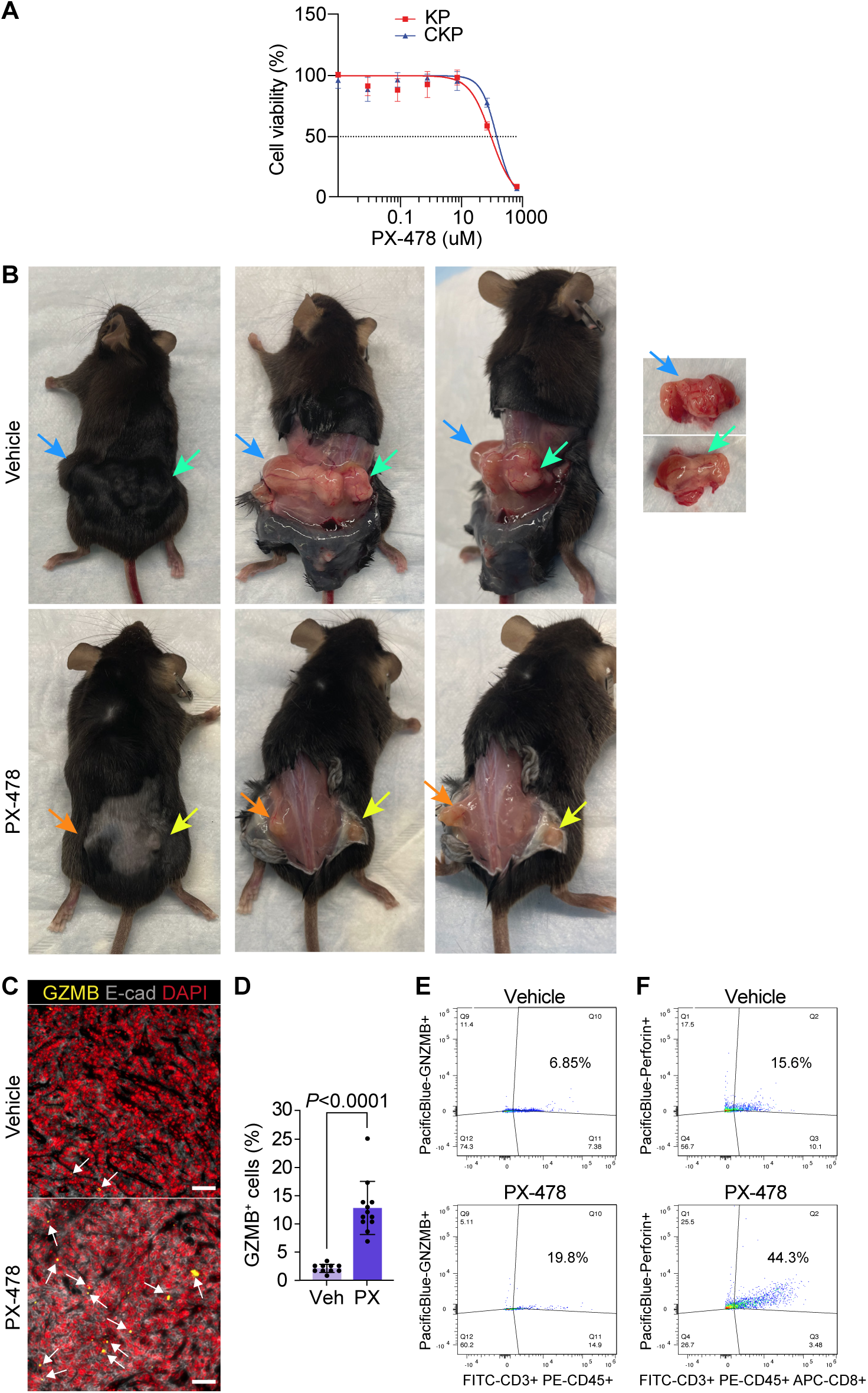

## References

1. Lin J-L, Lin J-X, Lin G-T, Huang C-M, Zheng C-H, Xie J-W, Wang J-b, Lu J, Chen Q-Y, Li P. Global incidence and mortality trends of gastric cancer and predicted mortality of gastric cancer by 2035. BMC public health. 2024;24(1):1763.

2. Smyth EC, Nilsson M, Grabsch HI, van Grieken NC, Lordick F. Gastric cancer. Lancet. 2020;396(10251):635–48. doi: 10.1016/S0140-6736(20)31288-5. PubMed PMID: 32861308.

3. Gao JP, Xu W, Liu WT, Yan M, Zhu ZG. Tumor heterogeneity of gastric cancer: From the perspective of tumor-initiating cell. World J Gastroenterol. 2018;24(24):2567–81. doi: 10.3748/wjg.v24.i24.2567. PubMed PMID: 29962814; PMCID: PMC6021770.

4. Oshi M, Satyananda V, Angarita FA, Kim TH, Tokumaru Y, Yan L, Matsuyama R, Endo I, Nagahashi M, Takabe K. Angiogenesis is associated with an attenuated tumor microenvironment, aggressive biology, and worse survival in gastric cancer patients. Am J Cancer Res. 2021;11(4):1659–71. Epub 20210415. PubMed PMID: 33948380; PMCID: PMC8085878.

5. Conti CB, Agnesi S, Scaravaglio M, Masseria P, Dinelli ME, Oldani M, Uggeri F. Early Gastric Cancer: Update on Prevention, Diagnosis and Treatment. Int J Environ Res Public Health. 2023;20(3). Epub 20230125. doi: 10.3390/ijerph20032149. PubMed PMID: 36767516; PMCID: PMC9916026.

6. Matsuoka T, Yashiro M. Novel biomarkers for early detection of gastric cancer. World J Gastroenterol. 2023;29(17):2515–33. doi: 10.3748/wjg.v29.i17.2515. PubMed PMID: 37213407; PMCID: PMC10198055.

7. Necula L, Matei L, Dragu D, Neagu AI, Mambet C, Nedeianu S, Bleotu C, Diaconu CC, Chivu-Economescu M. Recent advances in gastric cancer early diagnosis. World J Gastroenterol. 2019;25(17):2029–44. doi: 10.3748/wjg.v25.i17.2029. PubMed PMID: 31114131; PMCID: PMC6506585.

8. Alsina M, Arrazubi V, Diez M, Tabernero J. Current developments in gastric cancer: from molecular profiling to treatment strategy. Nat Rev Gastroenterol Hepatol. 2023;20(3):155–70. Epub 20221107. doi: 10.1038/s41575-022-00703-w. PubMed PMID: 36344677.

9. Chong IY, Chau I. Is there still a place for radiotherapy in gastric cancer? Curr Opin Pharmacol. 2023;68:102325. Epub 20230105. doi: 10.1016/j.coph.2022.102325. PubMed PMID: 36610101.

10. Joshi SS, Badgwell BD. Current treatment and recent progress in gastric cancer. CA Cancer J Clin. 2021;71(3):264–79. Epub 20210216. doi: 10.3322/caac.21657. PubMed PMID: 33592120; PMCID: PMC9927927.

11. Li K, Zhang A, Li X, Zhang H, Zhao L. Advances in clinical immunotherapy for gastric cancer. Biochim Biophys Acta Rev Cancer. 2021;1876(2):188615. Epub 20210814. doi: 10.1016/j.bbcan.2021.188615. PubMed PMID: 34403771.

12. Nakamura Y, Kawazoe A, Lordick F, Janjigian YY, Shitara K. Biomarker-targeted therapies for advanced-stage gastric and gastro-oesophageal junction cancers: an emerging paradigm. Nat Rev Clin Oncol. 2021;18(8):473–87. Epub 20210331. doi: 10.1038/s41571-021-00492-2. PubMed PMID: 33790428.

13. Lei ZN, Teng QX, Tian Q, Chen W, Xie Y, Wu K, Zeng Q, Zeng L, Pan Y, Chen ZS, He Y. Signaling pathways and therapeutic interventions in gastric cancer. Signal Transduct Target Ther. 2022;7(1):358. Epub 20221008. doi: 10.1038/s41392-022-01190-w. PubMed PMID: 36209270; PMCID: PMC9547882.

14. Sexton RE, Al Hallak MN, Diab M, Azmi AS. Gastric cancer: a comprehensive review of current and future treatment strategies. Cancer Metastasis Rev. 2020;39(4):1179–203. Epub 20200907. doi: 10.1007/s10555-020-09925-3. PubMed PMID: 32894370; PMCID: PMC7680370.

15. Lundquist MR, Storaska AJ, Liu T-C, Larsen SD, Evans T, Neubig RR, Jaffrey SR. Redox modification of nuclear actin by MICAL-2 regulates SRF signaling. Cell. 2014;156(3):563–76.

16. Grintsevich EE, Yesilyurt HG, Rich SK, Hung R-J, Terman JR, Reisler E. F-actin dismantling through a redox-driven synergy between Mical and cofilin. Nature cell biology. 2016;18(8):876–85.

17. Bear JE, Svitkina TM, Krause M, Schafer DA, Loureiro JJ, Strasser GA, Maly IV, Chaga OY, Cooper JA, Borisy GG. Antagonism between Ena/VASP proteins and actin filament capping regulates fibroblast motility. Cell. 2002;109(4):509–21.

18. Steffen A, Faix J, Resch GP, Linkner J, Wehland J, Small JV, Rottner K, Stradal TE. Filopodia formation in the absence of functional WAVE-and Arp2/3-complexes. Molecular biology of the cell. 2006;17(6):2581–91.

19. Biber G, Ben-Shmuel A, Sabag B, Barda-Saad M. Actin regulators in cancer progression and metastases: From structure and function to cytoskeletal dynamics. International Review of Cell and Molecular Biology. 2020;356:131–96.

20. Ockfen E, Filali L, Pereira Fernandes D, Hoffmann C, Thomas C. Actin cytoskeleton remodeling at the cancer cell side of the immunological synapse: good, bad, or both? Frontiers in Immunology. 2023;14:1276602.

21. Rouyère C, Serrano T, Frémont S, Echard A. Oxidation and reduction of actin: Origin, impact in vitro and functional consequences in vivo. European journal of cell biology. 2022;101(3):151249.

22. Zhang S, Kim K-B, Huang Y, Kim D-W, Kim B, Ko K-P, Zou G, Zhang J, Jun S, Kirk NA. CRACD loss promotes small cell lung cancer tumorigenesis via EZH2-mediated immune evasion. bioRxiv. 2023.

23. Jung Y-S, Wang W, Jun S, Zhang J, Srivastava M, Kim MJ, Lien EM, Shang J, Chen J, McCrea PD. Deregulation of CRAD-controlled cytoskeleton initiates mucinous colorectal cancer via β-catenin. Nature cell biology. 2018;20(11):1303–14.

24. Chang Y, Zhang J, Huo X, Qu X, Xia C, Huang K, Xie F, Wang N, Wei X, Jia Q. Substrate rigidity dictates colorectal tumorigenic cell stemness and metastasis via CRAD-dependent mechanotransduction. Cell Reports. 2022;38(7).

25. Kim B, Zhang S, Huang Y, Ko K-P, Jung Y-S, Jang J, Zou G, Zhang J, Jun S, Kim K-B. CRACD loss induces neuroendocrine cell plasticity of lung adenocarcinoma. Cell Reports. 2024;43(6).

26. Jung YS, Wang W, Jun S, Zhang J, Srivastava M, Kim MJ, Lien EM, Shang J, Chen J, McCrea PD, Zhang S, Park JI. Deregulation of CRAD-controlled cytoskeleton initiates mucinous colorectal cancer via beta-catenin. Nat Cell Biol. 2018;20(11):1303–14. Epub 20181022. doi: 10.1038/s41556-018-0215-z. PubMed PMID: 30361697; PMCID: PMC6261439.

27. Cristescu R, Lee J, Nebozhyn M, Kim KM, Ting JC, Wong SS, Liu J, Yue YG, Wang J, Yu K, Ye XS, Do IG, Liu S, Gong L, Fu J, Jin JG, Choi MG, Sohn TS, Lee JH, Bae JM, Kim ST, Park SH, Sohn I, Jung SH, Tan P, Chen R, Hardwick J, Kang WK, Ayers M, Hongyue D, Reinhard C, Loboda A, Kim S, Aggarwal A. Molecular analysis of gastric cancer identifies subtypes associated with distinct clinical outcomes. Nat Med. 2015;21(5):449–56. Epub 2015/04/22. doi: 10.1038/nm.3850. PubMed PMID: 25894828.

28. Cancer Genome Atlas Research N. Comprehensive molecular characterization of gastric adenocarcinoma. Nature. 2014;513(7517):202-9. Epub 20140723. doi: 10.1038/nature13480. PubMed PMID: 25079317; PMCID: PMC4170219.

29. Zou G, Huang Y, Zhang S, Ko KP, Kim B, Zhang J, Venkatesan V, Pizzi MP, Fan Y, Jun S, Niu N, Wang H, Song S, Ajani JA, Park JI. E-cadherin loss drives diffuse-type gastric tumorigenesis via EZH2-mediated reprogramming. J Exp Med. 2024;221(4). Epub 20240227. doi: 10.1084/jem.20230561. PubMed PMID: 38411616; PMCID: PMC10899090.

30. Ko KP, Huang Y, Zhang S, Zou G, Kim B, Zhang J, Jun S, Martin C, Dunbar KJ, Efe G, Rustgi AK, Nakagawa H, Park JI. Key Genetic Determinants Driving Esophageal Squamous Cell Carcinoma Initiation and Immune Evasion. Gastroenterology. 2023;165(3):613–28 e20. Epub 20230529. doi: 10.1053/j.gastro.2023.05.030. PubMed PMID: 37257519; PMCID: PMC10527250.

31. Ko KP, Zhang J, Park JI. Establishing transgenic murine esophageal organoids. STAR Protoc. 2022;3(2):101317. Epub 20220421. doi: 10.1016/j.xpro.2022.101317. PubMed PMID: 35496812; PMCID: PMC9048136.

32. Sun D, Guan X, Moran AE, Wu LY, Qian DZ, Schedin P, Dai MS, Danilov AV, Alumkal JJ, Adey AC, Spellman PT, Xia Z. Identifying phenotype-associated subpopulations by integrating bulk and single-cell sequencing data. Nat Biotechnol. 2022;40(4):527–38. Epub 2021/11/13. doi: 10.1038/s41587-021-01091-3. PubMed PMID: 34764492; PMCID: PMC9010342.

33. Moon KR, van Dijk D, Wang Z, Gigante S, Burkhardt DB, Chen WS, Yim K, Elzen AVD, Hirn MJ, Coifman RR, Ivanova NB, Wolf G, Krishnaswamy S. Visualizing structure and transitions in high-dimensional biological data. Nat Biotechnol. 2019;37(12):1482–92. Epub 20191203. doi: 10.1038/s41587-019-0336-3. PubMed PMID: 31796933; PMCID: PMC7073148.

34. Qin X, Rodriguez FC, Sufi J, Vlckova P, Claus J, Tape CJ. An oncogenic phenoscape of colonic stem cell polarization. Cell. 2023;186(25):5554–68. e18.

35. Davies A, Zoubeidi A, Beltran H, Selth LA. The Transcriptional and Epigenetic Landscape of Cancer Cell Lineage Plasticity. Cancer Discov. 2023;13(8):1771–88. doi: 10.1158/2159-8290.CD-23-0225. PubMed PMID: 37470668; PMCID: PMC10527883.

36. Van de Sande B, Flerin C, Davie K, De Waegeneer M, Hulselmans G, Aibar S, Seurinck R, Saelens W, Cannoodt R, Rouchon Q, Verbeiren T, De Maeyer D, Reumers J, Saeys Y, Aerts S. A scalable SCENIC workflow for single-cell gene regulatory network analysis. Nat Protoc. 2020;15(7):2247–76. Epub 20200619. doi: 10.1038/s41596-020-0336-2. PubMed PMID: 32561888.

37. Bonello S, Zahringer C, BelAiba RS, Djordjevic T, Hess J, Michiels C, Kietzmann T, Gorlach A. Reactive oxygen species activate the HIF-1alpha promoter via a functional NFkappaB site. Arterioscler Thromb Vasc Biol. 2007;27(4):755–61. Epub 20070201. doi: 10.1161/01.ATV.0000258979.92828.bc. PubMed PMID: 17272744.

38. Koshikawa N, Hayashi J, Nakagawara A, Takenaga K. Reactive oxygen species-generating mitochondrial DNA mutation up-regulates hypoxia-inducible factor-1alpha gene transcription via phosphatidylinositol 3-kinase-Akt/protein kinase C/histone deacetylase pathway. J Biol Chem. 2009;284(48):33185–94. Epub 20091001. doi: 10.1074/jbc.M109.054221. PubMed PMID: 19801684; PMCID: PMC2785161.

39. Page EL, Robitaille GA, Pouyssegur J, Richard DE. Induction of hypoxia-inducible factor-1alpha by transcriptional and translational mechanisms. J Biol Chem. 2002;277(50):48403–9. Epub 20021011. doi: 10.1074/jbc.M209114200. PubMed PMID: 12379645.

40. Bailey CM, Liu Y, Liu M, Du X, Devenport M, Zheng P, Liu Y, Wang Y. Targeting HIF-1alpha abrogates PD-L1-mediated immune evasion in tumor microenvironment but promotes tolerance in normal tissues. J Clin Invest. 2022;132(9). doi: 10.1172/JCI150846. PubMed PMID: 35239514; PMCID: PMC9057613.

41. Noman MZ, Desantis G, Janji B, Hasmim M, Karray S, Dessen P, Bronte V, Chouaib S. PD-L1 is a novel direct target of HIF-1alpha, and its blockade under hypoxia enhanced MDSC-mediated T cell activation. J Exp Med. 2014;211(5):781–90. Epub 20140428. doi: 10.1084/jem.20131916. PubMed PMID: 24778419; PMCID: PMC4010891.

42. Kakiuchi M, Nishizawa T, Ueda H, Gotoh K, Tanaka A, Hayashi A, Yamamoto S, Tatsuno K, Katoh H, Watanabe Y. Recurrent gain-of-function mutations of RHOA in diffuse-type gastric carcinoma. Nature genetics. 2014;46(6):583–7.

43. Kwon CH, Kim YK, Lee S, Kim A, Park HJ, Choi Y, Won YJ, Park DY, Lauwers GY. Gastric poorly cohesive carcinoma: a correlative study of mutational signatures and prognostic significance based on histopathological subtypes. Histopathology. 2018;72(4):556–68.

44. Koemans W, Luijten J, van der Kaaij R, Grootscholten C, Snaebjornsson P, Verhoeven R, van Sandick J. The metastatic pattern of intestinal and diffuse type gastric carcinoma–A Dutch national cohort study. Cancer epidemiology. 2020;69:101846.

