## Supplementary material for "Actin dysregulation induces immune evasion via oxidative stress-activated PD-L1 in gastric cancer": Methods

### Histology

#### Immunohistochemistry and immunofluorescence staining

The GC tumor microarray was purchased from Biomax (ST821). The murine and human tissue/tumor samples were fixed with 10% formalin, followed by paraffin embedding to make FFPE (formalin-fixed paraffin-embedding) blocks. The FFPE samples were immuno-stained following the standard protocol: deparaffinization, antigen retrieval, blocking, and primary antibody incubation. For immunohistochemistry (IHC) staining, HRP-conjugated secondary antibody was used. 3,3'-Diaminobenzidine (DAB) staining was performed with the ImmPACT DAB kit (Vector, SK-4105). The nuclei were counterstained with Hematoxylin QS Counterstain solution (Vector, H-3404-100). For the immunofluorescence staining, the fluorescence-conjugated secondary antibody was used. For immunofluorescence staining of cells, cells were cultured on the coverslip, followed by PBS wash (three times) and 4% paraformaldehyde solution fixation (15 min at room temperature). The nuclei were counterstained with ProLong Gold Antifade Mountant with DAPI (Thermo Fisher, P36935). Coverslips were mounted with ProLong Gold Antifade Mountant with DAPI and imaged. For the antibody information, see Table S1. Additional detals about TMA are provided in Table S2.

#### AB-PAS and HE staining

Deparaffinized tissue/tumor samples or paraformaldehyde (Fisher, #30525-89-4) fixed cells were incubated for 15 min with alcian blue (Sigma-Aldrich, A5268) solution (pH 2.5), followed by periodic acid-Schiff (PAS) staining (Sigma-Aldrich, 395B) according to the manufacturer's protocol. The nuclei were counterstained with Hematoxylin QS Counterstain solution or Nuclear Fast Red Counterstain solution (Vector, H-3403-500). For hematoxylin and eosin (HE) staining, the standard procedure was employed using HE staining kit (Abcam, ab245880).

#### Phalloidin slide staining

Tissue slides prepared using optimal cutting temperature (O.C.T) compound were either stored at 4°C or stained directly. Fifty microliters of Phalloidin was applied to the circled area and incubated at room temperature for 1 hour in the dark. Slides were washed in 0.1% PBST (Triton X-100 in PBS) three times for 5 minutes each. Sections were mounted using ProLong Gold Antifade Mountant with DAPI.

#### ROS detection

Cells were stained using the DCFDA - Cellular ROS Assay Kit (Abcam, ab113851) following the manufacturer's protocol. Briefly, cells were seeded in a 48-well and allowed to adhere overnight. The culture medium was removed, and the cells were washed with 1× PBS. Next, the cells were incubated with a working solution of 25 $\mu$M DCFDA in serum-free medium for 30 minutes at 37°C in the dark. After incubation, the cells were washed with 1× PBS, and fluorescence was measured using a microplate reader at excitation/emission wavelengths of 485 nm/535 nm to quantify ROS levels.

#### Microscopy

Images were captured by using Axio Observer Z1M (Zeiss), confocal (Nikon), and ECHO Revolve (ECHO), microscopes. Image adjustment was performed using ZEN software (v3.0, blue edition, Zeiss).

### RT-qPCR

RNAs were extracted by TRIzol solution (Invitrogen, #15596018) and used to synthesize cDNAs using the iScript Reverse Transcription Supermix (Bio-Rad, #1708841) following the manufacturer's protocols. RT-qPCR was performed using the amfiSure qGreen Q-PCR Master Mix (GenDEPOT, Q5602) at the Applied Biosystems 7500 Real-Time PCR System (ThermoFisher) with primers. Mouse *Cd274* (forward: 5'-GACGCAGGCGTTTACTGCT-3'; reverse: 5'- GCGGTATGGGGCATTGACTTT-3'). The comparative $2^{-\Delta\Delta Ct}$ method was used to quantify the fold changes of genes.

### Cell proliferation and viability assays

Cells were counted using a hemocytometer (Bio-Rad) on designated growth days, following the manufacturer's instructions. Cell proliferation was assessed either through crystal violet staining or by utilizing the Cell Counting Kit-8 (Dojindo Laboratories), in accordance with the manufacturer's protocol.

### Crystal violet staining

Cells (1 x $10^3$) were plated in 6-well plates, with the culture medium refreshed every 2 days. Afterward, the plates were washed with 1× PBS, fixed using 4% paraformaldehyde for 20 minutes, and stained with a crystal violet solution (0.1% crystal violet in 10% methanol) for 20 minutes. Excessive stains were removed by rinsing the plates with tap water.

### Immunoblot

Cell lysates were prepared by incubating the cell plate or the mashed tumor tissues with RIPA lysis buffer (50 mM Tris-HCl pH=7.5, 150 mM NaCl, 1.5 mM EDTA, 1% NP-40, 0.5% Na-deoxycholate, 0.1% SDS) for 30 min followed by full speed centrifugation for 10 min at 4°C. The protein concentration of the supernatant was calculated with a Pierce BCA protein assay kit (Thermo Fisher, #23227). The protein concentrations of loading samples were balanced with 5´ SDS loading buffer (2.5 ml 1 M Tris-HCl pH=6.8, 1 ml water, 1 g SDS powder, 5 ml glycerol, 0.5 ml β-mercaptoethanol, 0.5% bromophenol blue) and RIPA buffer, followed by denaturation at 95°C for 5 min. Then, the samples were loaded for SDS-PAGE. After the electrophoresis, the gel was transferred to the nitrocellulose membrane. The membrane was blocked with 5% non-fat dry milk in TBST (6.06 g Tris base powder, 8.76 g NaCl, 0.5 ml Tween-20 in 1 L water, pH=7.6) for 30 min at room temperature. Then, it was incubated with the primary antibody diluted in 2% BSA in TBST overnight at 4°C. Three times of washes with TBST were conducted. The HRP-conjugated secondary antibody was incubated with the membrane in TBST for 1 h at room temperature. Three times of washes with TBST were conducted. Enhanced chemiluminescence was performed with SuperSignal West Pico PLUS (Thermo Fisher, #34577) or SuperSignal West Femto Maximum Sensitivity Substrate (Thermo Fisher, #34095) reagent. For the antibody information, see Table S1.

### Flow cytometry

Tumors from syngeneic models were collected and processed into single-cell suspensions for flow cytometry analysis. Tumor tissues were minced with a blade and transferred into a digestion solution containing collagenase A and DNase I (Sigma). The tissue suspension was incubated at 37°C for 30 minutes to facilitate enzymatic digestion. After incubation, the cell suspension was filtered through a 70 µm cell strainer (Falcon) and washed twice with PBS. The cells were then filtered through a FACS tube strainer (Falcon) and washed twice more with FACS buffer (PBS containing 0.5% BSA and 2 mM EDTA). For surface staining, the following antibodies were used: PE anti-mouse CD45 (BioLegend, 1:100 dilution), Pacific Blue anti-mouse CD4 (BioLegend, 1:100 dilution), FITC anti-mouse CD3 (BioLegend, 1:50 dilution), and APC anti-mouse CD8 (BioLegend, 1:50 dilution). Cells were incubated with these antibodies for 30 minutes at 4°C in the dark. After staining, cells were washed twice with FACS buffer. For intracellular staining of perforin and Granzyme B, cells were fixed and permeabilized using the Fixation Solution (BioLegend)/Permeabilization Solution (3% TritonX100 in FACS buffer). Briefly, cells were fixed in fixation buffer for 20 minutes at 4°C, followed by washing with Perm/Wash buffer. Cells were then incubated with Pacific Blue anti-mouse perforin (BioLegend, 1:50 dilution) or Pacific Blue anti-mouse Granzyme B (BioLegend, 1:50 dilution) in Perm/Wash buffer for 1 hour at 4°C in the dark. After staining, cells were washed twice with Perm/Wash buffer and resuspended in FACS buffer. Flow cytometry was performed using an Attune flow cytometer, and data analysis was conducted with the FlowJo software.

### Animals

#### Syngeneic translpantation

C57BL/6 mice (6-7 weeks) were purchased from the Jackson Laboratory. Mice were randomized and subcutaneously injected with 3 x $10^6$ cells into both flanks. Mice were maintained in the Division of Laboratory Animal Resources facility at MD Anderson. Starting on day 7-10 after transplantation, mice were administered with PX-478 (40 mg/kg; intraperitoneal injection) or a-PD-L1 (10 mg/kg; intraperitoneal injection). Drug treatments were carried out approximately for 2-3 weeks, with administration every other day. a-PD-L1 antibody was injected twice a week for 4 weeks. Tumor volume was monitored and calculated by measuring with calipers every 2 days (volume = [length x width$^2$] / 2). Tumor burden was calculated by measuring all tumor lesions within the lung to account for the complete tumor burden. On day 25-30, mice were euthanized, tumors were photographed, and collected to proceed for paraffin-embedding and subsequent immunostaining or scRNA-seq.

#### *Cracd*$^{-/-}$ strain

The *Cracd*$^{-/-}$ mice (C57BL/6) were generated by the pronuclear injection of guide RNA (5'-TTCAT-GGGAA-TGGCG-TTCGA-3') integrated with an 80-mer SpCas9 scaffold into C57BL/6 mice by the Genetically Engineered Mouse Facility at MDACC[1].

#### KP strain

Compound transgenic mice *Kras*$^{LSL-G12D/+}$; *Trp53*$^{fl/fl}$ (KP) mice have been previously described[2]. C57BL/6 mice were purchased from The Jackson Laboratory

### *CRACD* mutation analysis

#### Mutation hotspots

*CRACD* mutation hotspot plots were produced by cBioPortal (www.cbioportal.org).

### Organoids

#### Gastric organoids generation and culture

Organoid models used in this study were previously established and characterized as described[3].

#### 2D culture

Organoids were dissociated following the organoid passaging protocol[3] and resuspended with RPMI1640 + 10% FBS with 10 *μ*M Y-27632. Then, organoids were seeded on a 24-well plate. Cells were passaged every 3-5 days. After the third passage, Y-27632 was removed from the culture medium.

#### FFPE sample preparation

Organoid pellet collected from PBS-flushed Matrigel, as described above, was further washed, and pelleted twice with cold PBS. Then, the organoids were fixed with 10% formalin and processed to the paraffin embedding.

#### Z-stack imaging and diameter measurement

Z-stack imaging of the Matrigel dome was performed under a 5x lens to measure the diameters of organoids. The diameters of organoids were quantified in pixels using ZEN software (Zeiss). The sizes were quantified in $\mu m^2$ with ImageJ[4] and converted to diameters in *μ*m.

### scRNA-seq

#### Single-cell isolation of GOs

For scRNA-seq, organoids from WT, C, KP, or CKP were collected 7 days after seeding and passed. Preparing GOs for scRNA-seq library preparation was performed as previously described[3].

#### Single-cell isolation of allograft tumors

Allograft tumors from mice (KP and CKP) were collected. The method for single-cell isolation of tumors was modified from the precvius study.[3] Tumors collected from mice were transferred to a 1.5 ml tube and dissected into small pieces (smaller than 2 mm) with scissors. Tumor pieces were then incubated with 1 ml tumor digestion buffer (2.5% FBS, 1 mg/ml Collagenase Type IV [STEMCELL, #07436], 250 µg/ml Dispase II [Gibco, #17105041], and 10 µM Y-27632 in advanced DMEM/F12 [advDF] medium [Gibco, #12634010]) at 37°C in a water bath for 30 min. After incubation, the tumor pieces were completely dissociated by robust pipetting with a p1000 pipette, followed by a resuspension with 4 ml cold PBS. Then, the digested mixture was passed through a 100 µm cell strainer to enrich single cells. The filtrate was pelleted by swing-bucket centrifugation under 300 g for 3 min at 4°C. Cells were further washed and pelleted twice by cold PBS. The supernatant was discarded. For FACS, the single-cell pellet was resuspended in cold 5% FBS-PBS. For organoid seeding, the pellet was resuspended in an appropriate volume of cold advDF plus medium. Then, the cell suspension was processed immediately for library preparation.

#### Library construction

scRNA-seq libraries were prepared using the CellPlex kit and the 10× Genomics Chromium Single Cell Gene Expression 3' v2 kit. A single-cell suspension was obtained by passing cells through a 35-µm cell strainer. Each group of cells was labeled with two Cell Multiplexing Oligo (CMO) tags from the CellPlex kit (10× Genomics). The tagged cells from different groups were pooled in equal proportions based on cell counts. Complementary DNA (cDNA) libraries were then constructed using the 10× Genomics 3' v2 kit. The libraries were sequenced on an Illumina NovaSeq platform (Novogene), and the resulting sequencing reads were aligned to the GRCm38/mm10 genome and demultiplexed using CellRanger (version 7.0.1). The generated count matrices were analyzed using Seurat (version 4.0.3) in R and Scanpy (version 1.8.2) in Python[5].

**Sequencing**

The libraries were sequenced using the Illumina NovaSeq platform at Novogene USA, with a read length of 150 paired-end base pairs and a sequencing depth of 200 million read pairs. The original sequencing data from the NovaSeq platform was transformed into raw reads by base calling. Raw reads were stored as fastq format files. Then, fastq files were processed to the data processing workflow.

**Data processing of raw sequencing reads and cell clustering**

Cell Ranger (v6.1.2, 10x Genomics) was used to align the raw sequencing reads to the reference genomes (mm10 for mouse, hg38 for human) to generate the 10x matrices. The ambient RNA and doublets of the 10x matrices were removed by SoupX (v1.6.2)[6] and Scrublet (v0.2.3)[7], respectively. Then, Scanpy (v1.8.2)[8] was used for processing the 10x matrices. Cells expressing less than 100 genes and genes expressed in less than 25 cells were filtered out. For mice, cells with less than 8000 counts per cell were kept. For humans, cells with less than 13000 counts per cell were kept. Cells expressing less than 50% mitochondrial genes and 50% ribosomal genes were kept. Then, filtered scRNA-seq data was normalized to 10,000 reads per cell and logarithmized. The top 5,000 highly variable genes were identified. The effects of total counts per cell and the percentages of mitochondrial and ribosomal genes were regressed out before scaling the data to unit variance. After the above procedures (called 'data quality control'), the dimensionality of the data was reduced by principal component analysis (PCA). For dataset integration, harmonypy (v0.0.5)[9] was performed to replace the PCA results (sce.pp.harmony_integrate(), basis='X_pca', adjusted_basis='X_pca_harmony'). The neighborhood graph of cells was computed (sc.pp.neighbors(), n_neighbors=15, n_pcs=50) using the PCA representation of the data matrix, clustered by the Leiden graph-clustering method (sc.tl.leiden(), resolution=0.5-1)[10], and embedded using UMAP (sc.pl.umap(), n_components=2 for 2D visualization, n_components=3 for 3D)[11]. The subset of *EPCAM+/Epcam+* cells was re-clustered following the same procedures as described above. Cell clusters were annotated according to the highly differential genes (i.e., marker genes) of each cluster determined by sc.tl.rank_genes_groups() (method='wilcoxon' for mouse, method='t-test' for human, use_raw=False).

### Cell plastic potential computation

The computation of cell plastic potential followed the protocol from Qin et al.[12]

**Single-cell entropies**

The single-cell entropy was computed by SCENT (v1.0.3)[13]. The normalized and logarithmized scRNA-seq data were converted from Scanpy to Seurat (v4.4.0) object[14]. The mouse gene symbols were converted to human Entrez Gene identifiers by Orthology.eg.db (v3.17.0) and org.Mm.eg.db (v3.17.0). The single-cell entropy was computed using CCAT (Correlation of Connectome And Transcriptome) algorithm (CompCCAT(), ppiA=net17Jan16.m). Then, the CCAT values of single cells were exported.

**RNA velocity lengths**

The RNA velocity lengths of single cells were exported from scVelo dynamical modeling as described above.

**Single-cell PHATE coordinates**

The PHATE embedding was computed by PHATE python package (v1.0.11)[15] using the normalized and logarithmized scRNA-seq data (phate_operator.fit_transform(adata.raw.X)). Then, the PHATE coordinates of single cells were exported.

**Valley-Ridge (VR) scores:** The VR score was defined as the weighted sum of the Valley (weight=0.9) and the Ridge (weight=0.1) components and was computed on a per sample and per cluster basis. The Valley component equals the median CCAT value of each sample-cluster combination. To calculate the Ridge component, the inverse of the RNA velocity length was first computed and scaled to a range between 0 and 1. The cell centrality distance was then calculated for cells in each cluster from the single-cell PHATE coordinates using Qin et al. [12] defined Python function (compute_distdeg(), knn value was optimized according to the size of each cluster). Finally, the Ridge component was computed per sample-cluster as the product of the median scaled inverse velocities and the cell's scaled centrality distance.

##### Waddington-like landscapes

The Waddington-like landscapes were visualized in Houdini Indie (SideFX, v20.0.533). The VR scores were positioned on the y-axis and the single-cell PHATE coordinates were positioned on the xz plane.

#### Cell-cell communications

For the analysis of ligand-receptor interaction-based cell-cell communication in single-cell RNA sequencing (scRNA-seq) datasets, the CellChat package in R (https://www.r-project.org) was utilized. The integrated dataset was preprocessed using the Seurat package and subsequently clustered and annotated. The processed dataset was then analyzed using CellChat with default parameters, including a *P* value threshold of 0.05. Epithelial cells were designated as the source group, while immune cells served as the target group.

#### Regulatory network inference

First, cells expressing less than 100 genes and genes expressed in less than 25 cells were filtered out. Cells with less than 8000 counts/cell and cells expressing less than 50% mitochondrial genes and 50% ribosomal genes were kept. Second, given a manually curated list of 1721 murine TFs, regulatory interactions between these TFs and putative target genes were inferred via gradient boosting machine regression (GRNBoost2) from the filtered scRNA-seq data (pyscenic grn). Multiple modules of different sizes were generated from these regulatory interactions. Each module consisted of a TF and its predicted target genes. Third, enriched motifs in the regulatory regions of target genes were identified by comparing cis-regulatory module scores near these genes. The AUC metric was used to assess significant gene recovery within a whole genome ranking. All direct target genes from modules sharing a regulator were compiled into one final regulon (pyscenic ctx). Fourth, the AUC metric measured the relative biological activity of the predicted regulons in the individual cells of the scRNA-seq data (pyscenic aucell). Finally, the activity of the distribution of the cellular AUC values for regulons was binarized by binarize() and plotted as a heatmap. The UMAP of CA tumor annotated by regulon patterns was re-clustered by AUCell (sc.pp.neighbors(), n_neighbors=15, n_pcs=50, use_rep='X_aucell'). RSSs were computed by regulon_specificity_scores(). The feature plots visualizing the TF expression localizations and regulon activity localizations were plotted by sc.pl.umap().

#### Cell cycle scoring

To determine the cell cycle states of individual cells, we utilized the sc.tl.score_genes_cell_cycle function from the Scanpy package. This method calculates cell cycle scores for each cell based on the expression of phase-specific marker genes [PMID: 27124452], allowing for the classification of cells into distinct cell cycle states (G1, S, and G2/M).

#### Pseudotime analysis using CytoTRACE

Pseudotime trajectories were inferred using the CytoTRACE R package (v 0.3.3), which estimates the differentiation state of single cells based on their transcriptional diversity. The algorithm assigns a differentiation score to each cell, where higher scores indicate greater transcriptional diversity and a less differentiated state. These scores were subsequently used to order cells along a pseudotime trajectory. CytoTRACE-generated pseudotime values were visualized using UMAP and violin plots to illustrate the progression of cellular states.

### GSVA and GSEA

#### Data downloading of TCGA-STAD

The bulk RNA-seq data of TCGA-STAD patients was prepared by TCGAbiolinks. Briefly, the count matrix was downloaded by GDCquery(project="TCGA-STAD", data.category="Transcriptome Profiling", data.type="Gene Expression Quantification", workflow.type="STAR - Counts", experimental.strategy='RNA-Seq', sample.type=c('Primary Tumor', 'Solid Tissue Normal'). The clinical data was downloaded by GDCquery_clinic(project="TCGA-STAD", type="Clinical"). Based on the sample barcodes (01 for 'Solid Tissue Normal', 11 for 'Primary Solid Tumor') and the 'histological_type' (only 'Stomach, Adenocarcinoma, Diffuse Type' was included) in the clinical data, the count matrix was sorted in the order of Normal, and diffuse type STAD. The human gene Ensembl IDs inside of the count matrix were converted to HUGO gene symbols by biomaRt.

### ATAC-seq

#### Single-cell isolation

The single-cell isolation was performed as described above in the section 'Organoids'. Two individual biological replicates per group were used. The final single-cell pellet was resuspended in 5% FBS-PBS and proceeded to FACS immediately.

#### FACS for live EPCAM$^+$ cells

The single-cell pellet prepared from tumors was first resuspended in 100 µl 5% FBS-PBS with 1 µl PE/Cyanine7-Epcam antibody and incubated on ice for 30 min. Afterward, cells were washed once with 10 ml cold 5% FBS-PBS and pelleted by swing-bucket centrifugation under 300 g for 3 min at 4°C. The supernatant was discarded. Then, the pellet was resuspended in 100 µl 5% FBS-PBS with 5 µl 7-AAD Viability Staining Solution (Invitrogen, #00-6993-50) and incubated on ice for 5 min. After the incubation, the cell suspension was diluted to 500 µl per group and proceeded to FACS immediately. The cell suspension was loaded onto the FACSAria II cell sorter (BD). EPCAM$^+$ & 7AAD$^-$ cells were sorted out and resuspended in cold PBS for the library construction immediately. For the antibody information, see Table S24.

#### Library construction

The ATAC-seq library construction method was modified from the Omni-ATAC-seq protocol[16]. Briefly, 75,000 viable cells after FACS per group were counted and pelleted by fixed-angle centrifugation under 400 g for 4 min at 4°C. The supernatant was carefully discarded. The cell membrane was disrupted by adding 50 µl cold ATAC-resuspension buffer (10 mM Tris-HCl pH=7.4, 10 mM NaCl, 3 mM $MgCl_2$ in nuclease-free water) containing 0.1% NP40 (Roche, 11332473001), 0.1% Tween-20, and 0.01% digitonin (Invitrogen, BN2006) with 3 times of pipetting. After the incubation on ice for 3 min, the lysis was washed out by adding 1 ml cold ATAC-resuspension buffer containing 0.1% Tween-20 only with 3 times of pipetting. The nuclei were pelleted by a fixed-angle centrifugation under 400 g for 8 min at 4°C. The supernatant was carefully discarded. The nuclei pellet was resuspended by 50 µl transposition mix (25 µl 2x TD buffer and 2.5 µl transposase from the Tagment DNA Enzyme and Buffer Small Kit [Illumina, #20034197], 16.5 µl PBS, 0.5 µl 1% digitonin, 0.5 µl 10% Tween-20, 5 µl nuclease-free water) with 6 times of pipetting. This reaction mixture was incubated at 37°C for 30 min in a thermomixer with 1000 rpm mixing. After the transposition, the reaction mixture was cleaned up with a DNA Clean & Concentrator-5 kit (ZYMO, D4013), followed by elution. The eluted mixture per group was pre-amplified for 5 cycles of PCR using different primers, respectively. Then, 5 µl of the pre-amplified mixture was used to run a 15 µl qPCR amplification to determine the number of additional cycles needed. Finally, the remainder of the pre-amplified DNA ran the additional PCR cycles. The amplified library was purified and eluted in 20 µl nuclease-free water. The libraries were subsequently subjected to sequencing.

#### Sequencing

The libraries were sequenced using the Illumina NovaSeq platform at Novogene USA, with a read length of 150 paired-end base pairs and a sequencing depth of 50 million read pairs. The original sequencing data from the NovaSeq platform was transformed into raw reads by base calling. Raw reads were stored in fastq format files. Then, the fastq files were processed for the bioinformatic analyses.

#### Motif-predicted binding sites

Bioinformatic analyses of ATAC-seq, including quality control of raw reads, adapter trimming, alignment with the reference genome, duplicate marking, reads filtering, peak calling, peak annotation, PCA, and replicates merging, were performed by nf-core/atacseq pipeline (v2.1.2)[17]. The insert size distribution plot, the heatmap of peaks binding to TSS regions, the Euclidean distance matrix, and peak annotation bar plot were generated by this pipeline. The merged bigwig file per group containing tracks and peaks was loaded in the integrative genomics viewer for visualization.

### CUT&RUN

#### CUT&RUN assays

CUTANA ChIC/CUT&RUN kit (EpiCypher, Cat. No. 14-1048) was used based on the manufacturers' protocols. CUT&RUN was performed using *HIF1a* antibodies (Cell Signaling, 14179 and 36169). For controls, IgG and H3K4me3 antibodies provided in the kit were used.

#### PCR and qRT-PCR

CUT&RUN was performed using *HIF1a* antibodies, and the resulting DNA was used to evaluate the binding of *HIF1a* to *Cd274*. *Cd274* primers (p1 F: 5'-TGAAGT GTCTGG ATTCTG AAG-3', R: 5'-TTAGGG TGACCT TTGGGATA-3'; p2 F: 5'-GAGGAA GTCACC AAATCCAC-3', R: 5'-TCTTGA AAGCCC TTTCTGGA-3'; p3 F: 5-ATCCAC GTATCC AGAAAGGG-3', R: 5'-TCGGTG GTGGTG ACTACTG-3'; p4 F: 5'-TCTTAT GACTTCAGATAT TTTGCT-3', R: 5'-AGGAGC AGTGAGCGC TTTAG-3'; p5 F: 5'-ATGACTGGGTCTTTCCACTT-3', R: 5'-GAGGTCTAGGATGCTGGAGC-3'; p6 F: AAACAGTTCTTAGAT ACAGTG-3', R: 5'-GGGTTGTCATAAGAGGTGAAA-3') were designed using the VISTA browser and used for PCR (Genedepot, PCR master mix) under the manufacturer's recommended conditions. qRT-PCR was subsequently performed, following the methods described earlier.

### Cell lines

#### Cell culture

2D organoid cell lines KP and CKP were cultured with RPMI1640 + 10% FBS + 1% PS at 37°C in a humidified incubator supplied with 5% $CO_2$, as we previously performed[3].

#### Drugs and Chemical testing on cell lines

PX-478 (Selleckchem, S7612) was obtained commercially. To determine GI50 values, cells were digested with Trypsin-EDTA (Corning, #25-052-CI) to prepare a single-cell suspension. The suspension was diluted to a concentration of 30,000 cells per ml, and 100 µl of this suspension was seeded into each well of a 96-well plate, leaving the last row blank without cells. After 24 hours of incubation, 200 µl of medium containing drugs at gradient concentrations was added to each well, with triplicates for each concentration. Following a 72-hour incubation period, the drug-containing medium was replaced with 100 µl of fresh medium containing 5 µl of CCK-8 reagent from the Cell Counting Kit-8 (Dojindo, CK0413) per well. The plates were incubated at 37°C for an additional 4 hours, and absorbance at 450 nm was measured using a microplate reader. For analysis, absorbance values from the blank wells were used as 0% viability, while those from drug-free wells were set as 100% viability. GI50 values were calculated using GraphPad Prism. Latrunculin A (Cayman, 10010630) and Cytochalasin D (MCE, HY-N6682) were used to inhibit actin polymerization. Diphenyleneiodonium chloride (MCE, HY-100965), N-Acetylcysteine amide (MCE, HY-110256), and mitoTEMPO (MCE, HY-112879) were utilized to inhibit reactive oxygen species (ROS).

### Statistical analysis

GraphPad Prism was used for statistical analysis. The statistical significance tests were performed as each section described. *P* values lower than 0.05 are considered statistically significant. Error bars indicate the standard deviation (SD). All experiments were performed three or more times independently under identical or similar conditions.

### Graphic illustration

Cartoons were created with BioRender.com and modified with Adobe Illustrator.

### Resource availability

#### Lead contact

**Materials availability**

All reagents and models, including genetically engineered mice and organoid models, will be available upon request.

**Data and code availability**

scRNA-seq data is available via the Gene Expression Omnibus (GEO; accession number: GSE284638; token for reviewers: ).

ATAC-seq data is available from the corresponding author, J.-I.P., upon request. The code used to reproduce the analyses described in this study can be accessed via Github (github.com/jaeilparklab/CRACD_DGAC).

### References (for Methods)


1 Jung, Y. S. *et al.* Deregulation of CRAD-controlled cytoskeleton initiates mucinous colorectal cancer via beta-catenin. *Nat Cell Biol* **20**, 1303-1314 (2018). https://doi.org:10.1038/s41556-018-0215-z

2 Kim, M. J. *et al.* PAF remodels the DREAM complex to bypass cell quiescence and promote lung tumorigenesis. *Molecular cell* **81**, 1698-1714. e1696 (2021).

3 Zou, G. *et al.* E-cadherin loss drives diffuse-type gastric tumorigenesis via EZH2-mediated reprogramming. *Journal of Experimental Medicine* **221**, e20230561 (2024).

4 Schneider, C. A., Rasband, W. S. & Eliceiri, K. W. NIH Image to ImageJ: 25 years of image analysis. *Nat Methods* **9**, 671-675 (2012). https://doi.org:10.1038/nmeth.2089

5 Ko, K.-P. *et al.* Key genetic determinants driving esophageal squamous cell carcinoma initiation and immune evasion. *Gastroenterology* **165**, 613-628. e620 (2023).

6 Young, M. D. & Behjati, S. SoupX removes ambient RNA contamination from droplet-based single-cell RNA sequencing data. *Gigascience* **9** (2020). https://doi.org:10.1093/gigascience/giaa151

7 Wolock, S. L., Lopez, R. & Klein, A. M. Scrublet: Computational Identification of Cell Doublets in Single-Cell Transcriptomic Data. *Cell Syst* **8**, 281-291 e289 (2019). https://doi.org:10.1016/j.cels.2018.11.005

8 Wolf, F. A., Angerer, P. & Theis, F. J. SCANPY: large-scale single-cell gene expression data analysis. *Genome Biology* **19**, 15 (2018). https://doi.org:10.1186/s13059-017-1382-0

9 Korsunsky, I. *et al.* Fast, sensitive and accurate integration of single-cell data with Harmony. *Nat Methods* **16**, 1289-1296 (2019). https://doi.org:10.1038/s41592-019-0619-0

10 Traag, V. A., Waltman, L. & van Eck, N. J. From Louvain to Leiden: guaranteeing well-connected communities. *Scientific reports* **9**, 5233 (2019). https://doi.org:10.1038/s41598-019-41695-z

11 Becht, E. *et al.* Dimensionality reduction for visualizing single-cell data using UMAP. *Nat Biotechnol* (2018). https://doi.org:10.1038/nbt.4314

12 Qin, X. *et al.* An oncogenic phenoscape of colonic stem cell polarization. *Cell* **186**, 5554-5568.e5518 (2023). https://doi.org:10.1016/j.cell.2023.11.004

13 Teschendorff, A. E. & Enver, T. Single-cell entropy for accurate estimation of differentiation potency from a cell's transcriptome. *Nature communications* **8** (2017). https://doi.org:10.1038/ncomms15599

14 Stuart, T. *et al.* Comprehensive Integration of Single-Cell Data. *Cell* **177**, 1888-1902.e1821 (2019). https://doi.org:10.1016/j.cell.2019.05.031

15 Moon, K. R. *et al.* Visualizing structure and transitions in high-dimensional biological data. *Nature Biotechnology* **37**, 1482-1492 (2019). https://doi.org:10.1038/s41587-019-0336-3

16 Corces, M. R. *et al.* An improved ATAC-seq protocol reduces background and enables interrogation of frozen tissues. *Nat Methods* **14**, 959-962 (2017). https://doi.org:10.1038/nmeth.4396

17 Ewels, P. A. *et al.* The nf-core framework for community-curated bioinformatics pipelines. *Nat Biotechnol* **38**, 276-278 (2020). https://doi.org:10.1038/s41587-020-0439-x
