## Supplementary Information for "Actin dysregulation induces immune evasion via oxidative stress-activated PD-L1 in gastric cancer"

#### Supplementary Figures

Supplementary Figure 1. Genetic alterations of *CRACD* in gastric cancer
Supplementary Figure 2. Analysis of *Cracd*-depleted gastric epithelia and organoids
Supplementary Figure 3. Single-cell transcriptomic analysis of GOs
Supplementary Figure 4. Analyses of allograft tumors
Supplementary Figure 5. Comparative analyses of gene regulatory networks
Supplementary Figure 6. Impact HIF1α inhibition on CKP tumorigenesis and immune modulation

#### Supplementary Tables (.xlsx file)

Supplementary Table 1. Reagent information
Supplementary Table 2. Tissue microarray information (ST821)

#### Uncropped images of immunoblots

### Supplementary Figures

**Supplementary Figure 1. Genetic alterations of *CRACD* in gastric cancer**
**A.** cBioPortal analysis of TCGA Pan-Cancer datasets. 10,967 samples; the plot from cancer type detailed. **B, C.** Immunohistochemistry (IHC) of gastric cancer tissue microarray for CRACD protein; hematoxylin for nuclear counterstaining. Scale bars, 100 μm. The representative images are shown (B) with quantitative analysis (C).

**Supplementary Figure 2. Analysis of Cracd-depleted gastric epithelia and organoids**
**A.** IHC f the mouse stomach tissues (*Cracd* WT or KO) for CRACD protein expression. **B.** Bright-field images of the stomach tissues isolated from *Cracd* WT or KO mice (7-month-old). Hyperplasia of the gastric epithelia was observed in *Cracd* KO mice. **C-H.** Histological analysis of the stomach tissues isolated from *Cracd* WT or KO mice. Hematoxylin and eosin (HE) (C), immunostaining for MKI67, a marker for proliferative cells (D), Alcian Blue–Periodic Acid–Schiff (AB-PAS), a marker for mucinous cells (E), phalloidin, a marker for filamentous actin (F), β-catenin (G), and E-cadherin (H). Yellow arrows point out the cells with increased β-catenin expression. Quantitation was performed using Student's *t*-test. **I-M.** Histological analysis of gastric organoids (GOs) derived from the gastric epithelial cells isolated from *Cracd* WT or KO mice. HE (I), MKI67 (J), Periodic Acid–Schiff (PAS) (K), and wholemount staining for phalloidin (L), E-cadherin (M). Quantitation was performed using Student's *t*-test and MKI67+ cells of WT, C, KP, and CKP GOs with one-way ANOVA. **N.** Genotyping of *Trp53*$^{fl/fl}$, KP, and CKP GOs for PCR with *Trp53*$^{fl/fl}$ and *Trp53*$^{del/del}$ primers. **O.** Genotyping of *Trp53*$^{fl/fl}$, KP, and CKP GOs with *Kras* WT and *Kras*$^{G12D}$ primers. **P.** Sanger sequencing results of KP and CKP GOs with *Cracd* primers. **Q.** Sanger sequencing results of KP and CKP (clone 1) GOs with *Cracd* primers. **R.** Bright-field images of *Trp53*$^{fl/fl}$, KP, and CKP GOs. **S.** Bromodeoxyuridine (BrdU) incorporation assay of *Trp53*$^{fl/fl}$, KP, and CKP GOs. **T.** Immunofluorescent (IF) staining of *Trp53*$^{fl/fl}$, KP, and CKP GOs for CEA5. Scale bars: green = 50 μm, blue = 10 μm; Student's *t*-test (unless specified); error bars: standard deviation; n>3; Representative images are shown; ns: not significant.

**Supplementary Figure 3. Single-cell transcriptomic analysis of GOs**
**A.** Dot plot showing the expression of genes related to gastric epithelium stemness in each type of GOs (WT, C, KP, and CKP). **B.** Feature plots of *Mki67*, *Sox4*, and *Aqp5* of WT, C, KP, and CKP GOs. **C, D.** Dot plot showing the expression of mucinous (C) and clinical (D) markers in each type of GOs. **E.** Feature plots displaying the expression of *Muc1* and *Krt7* of WT, C, KP, and CKP GOs.

**Supplementary Figure 4. Analyses of allograft tumors**
**A, B.** Statistical analysis of MKI67- (A) and CEA5- (B) positive cells in allograft tumors (KP vs. CKP). Error bars=SD; ***: $P < 0.001$, *: $P < 0.05$**.** **C, D.** Cell-cell interaction analysis of tumors (KP vs. CKP). Relative and absolute information flow of ligand-receptor interactions in KP and CKP tumor cells; analyzed using the CellChat package of scRNA-seq datasets of whole tumors (KP vs. CKP) (C) and **i**nteraction strength network between cell populations in KP and CKP tumors (D).

**Supplementary Figure 5. Comparative analyses of gene regulatory networks**
**A.** Feature plots visualizing the expression of transcription factor (TF), *Hif1a*, *Sox4*, and *Sox11* on the UMAPs of each GOs (WT, C, KP, and CKP). **B, C.** ATAC-seq analysis of KP and CKP tumor cells. Stacked bar plots displaying the distribution of TFs' binding loci relative to transcription start sites (TSS) (B) and features (C). **D.** Activity scores of transcription factors in KP and CKP tumors, showing differential regulatory networks. Highlighted transcription factors, including *Hif1a*, *Sox4*, and *Sox11*, exhibit higher activity in CKP cells. Red dots represent transcription factors with significant alterations between the two groups (KP and CKP).

**Supplementary Figure 6. Impact HIF1α inhibition on CKP tumorigenesis and immune modulation**
**A.** Cell viability assay of KP and CKP 2D cell lines treated with a *HIF1α* inhibitor (PX-478). **B.** Photos of immunocompetent mice (C57BL/6; vehicle vs. PX-478) bearing CKP allograft tumors. Tumors display invasive growth, penetrating the muscularis propria, and diffuse tumor formation. **C, D.** IF staining images (C) and quantification (D) of CKP allograft tumors post-*HIF1α* inhibitor treatment for Granzyme B (GZMB) and E-cadherin. Scale bars: 50 μm; arrows: Granzyme B and E-cadherin double-positive cells. **E, F.** Flow cytometric analysis of whole cells in CKP allograft tumors from mice treated with vehicle (control) or PX-478 for Granzyme B-positive

(E) or Perforin-positive (F) T cells, a *HIF1α* inhibitor. Student's *t*-test; error bars: standard deviation; n>3; Representative images and plots are shown.

**Uncropped images of immunoblots**

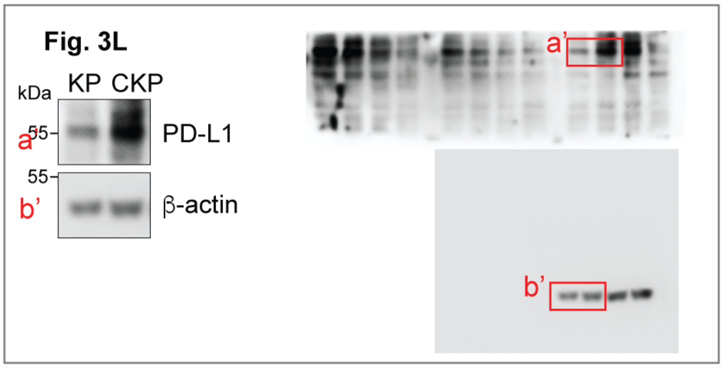


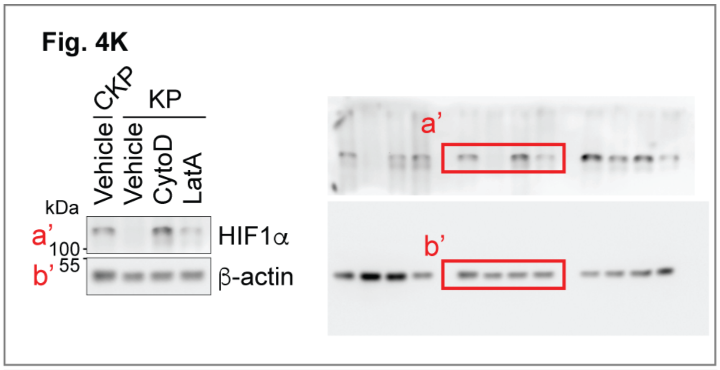


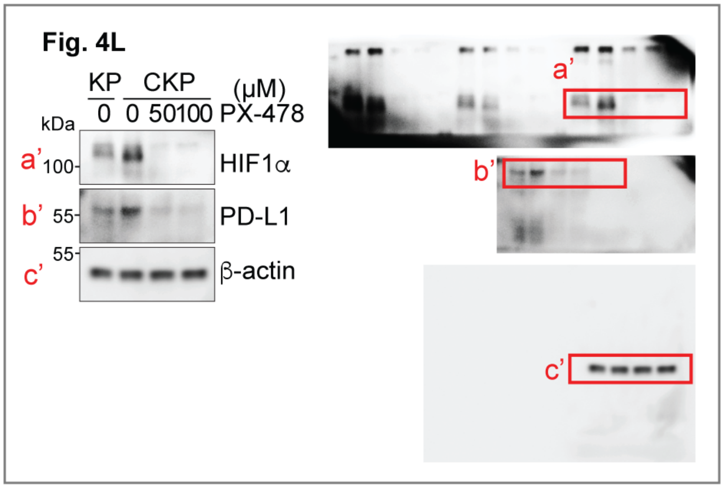


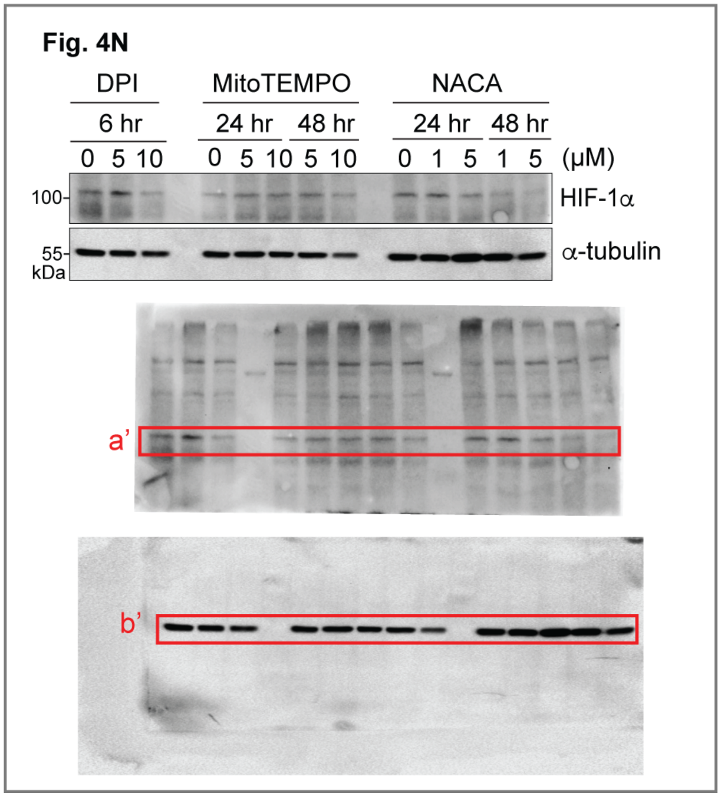
